# The inter-dimeric interface controls function and stability of *Ureaplasma urealiticum* methionine S-adenosyltransferase

**DOI:** 10.1101/669069

**Authors:** Daniel Kleiner, Fannia Shmulevich, Raz Zarivach, Anat Shahar, Michal Sharon, Gili Ben-Nissan, Shimon Bershtein

## Abstract

Methionine S-adenosyltransferases (MATs) are predominantly homotetramers, comprised of dimers of dimers. The highly conserved dimeric interface harbors two active sites, making the dimer the obligatory functional unit. Yet, functionality of the recently evolved inter-dimeric interface remains unknown. Here, we show that the inter-dimeric interface of *U. urealiticum* MAT has evolved to control the catalytic activity and structural integrity of the homotetramer in response to product accumulation. When all four active sites are occupied with the product, S-adenosylmethionine (SAM), binding of four additional SAM molecules to the inter-dimeric interface prompts a ∼45° shift in the dimer orientation and a concomitant ∼60% increase in the interface area. This rearrangement inhibits the enzymatic activity by locking the flexible active site loops in a closed state and renders the tetramer resistant to proteolytic degradation. Our findings suggest that the inter-dimeric interfaces of MATs are recruited by evolution to tune the molecular properties of the entire homotetramer.

## Introduction

S-adenosyltransferases (MATs) are highly conserved enzymes found in all kingdoms of life, predominantly as dihedral homotetramers (Markham and Pajares, 2009; Sanchez-Perez et al., 2003). Like other dihedral homotetramers, MAT comprises a dimer of dimers. MATs catalyze the only known biochemical route to formation of S-adenosylmethionine (SAM) ̶ a metabolite involved in myriads of metabolic reactions in the cell, including methylation, polyamine synthesis, and production of cofactors(Fontecave et al., 2004). The MAT monomer consists of three domains, each built of βαββαβ repeats (Markham and Pajares, 2009). To form a dimer, two monomers are paired in the inverted configuration, such that the α-helixes are surface exposed, whereas β-strands form a large flat hydrophobic interface. The interface is isologous, as the same surface patch from both subunits participates in its formation. Two deep cavities harboring active sites are located directly in this dimeric interface. Both monomers contribute to the formation of each of the active sites, making the homodimer the obligatory functional unit. Two such dimers the interact via a smaller isologous interface to form a homotetramer. Comparison of MAT structures from *E. coli (*eMAT) and rat liver (rlMAT) reveals that, unlike the large dimeric interfaces, the smaller inter-dimeric interfaces of these structures are highly diverged. A relatively small solvent-accessible interaction core in rlMAT interface is contrasted by a much more extended interface in eMAT (Gonzalez et al., 2000; Sanchez-Perez et al., 2003; Takusagawa et al., 1996). Whilst the emergence of the first dimeric interface is clearly adaptive, determining whether the assembly of two dimers into a tetramer is a functionally beneficial result of selection is not straightforward, particularly in the light of the fact that dimeric MATs have also been reported (Sanchez del Pino et al., 2002; Zano et al., 2014).

What are the mechanisms governing the emergence and assembly of dihedral homotetramers? An idea has been put forward suggesting that the evolutionary route towards assembly of dihedral homotetramers can be predicted based solely on the differences in the interface sizes (Levy et al., 2008). According to this idea, which is largely based on energetic considerations, the largest interface in the dihedral homotetramer has arose first in evolution via homodimerization of two monomers. The assembly of two dimers into a tetramer, mediated by the smaller interface, then followed. The hierarchy in the sizes of the interfaces, therefore, is preserved in evolution in order to maintain the proper assembly route. This idea is supported by the analytical predictions and experimental findings that show that most (if not all) dihedral homotetramers indeed follow the monomer-dimer-tetramer (MDT) (dis)assembly path (Levy et al., 2008; Powers and Powers, 2003; Villar et al., 2009).

Whilst the properties of the interfaces can teach us about the evolution of assembly of dihedral complexes, the functional contribution of the interfaces, and especially the recently evolved non-essential ones, is less obvious. Indeed, both adaptive and non-adaptive processes seem to contribute to the emergence of homomeric complexes (Marsh and Teichmann, 2015). Lynch has demonstrated a strong dependence of the pathways for the emergence of protein complexes on the effective population size of major phylogenetic groups and suggested that protein multimerization can be a stochastic outcome of random genetic drift (Lynch, 2012, 2013). Protein homomerization was also shown to emerge as a side effect to thermodynamic stabilization of the complexes, a process that does not necessarily lead to new or improved functionality (Bershtein et al., 2012; Fraser et al., 2016; Jacobs et al., 2016). Notwithstanding, probably all homomeric complexes possess the inherent potential for allosteric regulation (Fairman et al., 2011; Gunasekaran et al., 2004; Hilser et al., 2012) and functional activation (Hashimoto et al., 2011; Nooren and Thornton, 2003), phenomena that can eventually be recruited by adaptive evolution. Indeed, a strong relationship has been observed between symmetry of the homomeric complexes and function they perform, suggesting that protein function is an important determinant of quaternary structure evolution. Dihedral homomers, in particular, were found to be significantly enriched with allosteric metabolic enzymes (Bergendahl and Marsh, 2017).

Here we report a structure-function study of the yet uncharacterized S-adenosyltransferase (MAT) from *U. urealiticum* (uMAT). We show that the inter-dimeric interface plays a pivotal role in the regulation of enzymatic activity and structural integrity of uMAT homotetramer. Specifically, in response to an increase in SAM concentration, uMAT undergoes a dramatic structural rearrangement, whereby all active site loops attain a closed state facilitated by binding of SAM molecules to the inter-dimeric interface. Concomitantly, uMAT dimers rotate by ∼45%, thus increasing the surface of the inter-dimeric interface by 60%, and forming a continuous β-sheet that runs throughout the inter-dimeric interface and stiches both dimers together. This rearrangement inhibits the enzymatic catalysis and renders the tetramer resistant to proteolysis. We suggest that, unlike the highly conserved and evolutionary ancient large interface, the smaller and much more diverged inter-dimeric interface of MATs is recruited by evolution to tune the molecular properties of the entire homotetrameric complex. By acting on the smaller interface, evolution can tailor degradation propensity, activity regulation, and other crucial properties of the complex to the particular needs of the organism, whilst preserving the original function and assembly path dictated by the larger interface.

## Results

### uMAT is a dihedral homotetramer

We began by solving the structure of uMAT in the absence of ligands. Two virtually identical crystal structures, formed under different conditions and occupying different space groups (P1 and P21), were solved by homologous replacements, each to 2.6Å resolution (Figure 1A, Figure S1A) (Table S1) (**STAR Methods**). The overall structure of uMAT monomer is highly conserved: eMAT and uMAT monomers that share 44% sequence show root mean square deviation (RMSD) among α-carbons of 1.35Å (Figure S1B). uMAT is a dihedral homotetramer comprised of two isologous interfaces (Figure 1A,E,H, Table 1). The larger interface is formed between two monomers that interact along flat β-sheets to form a dimer (Figure 1E, Table 1, and Table S2). Apart from the numerous hydrophobic and H-bond interactions, the dimeric interface is also fortified by ∼12 salt-bridges. The calculated size of the interface (∼2,300Å^2^) is somewhat underestimated because of a few unresolved loop segments ̶ most notably a missing 13 amino acids-long fragment (positions 96-108) out of 28 amino acids-long flexible loop gating the access to the active site (positions 89-116) (Figure S1A, **STAR Methods**). The uMAT and eMAT dimers share a fairly similar topology (RMSD = 3.5Å), supporting the notion that homodimerization along the large interface is highly conserved (Figure S1C).

**Table 1.**
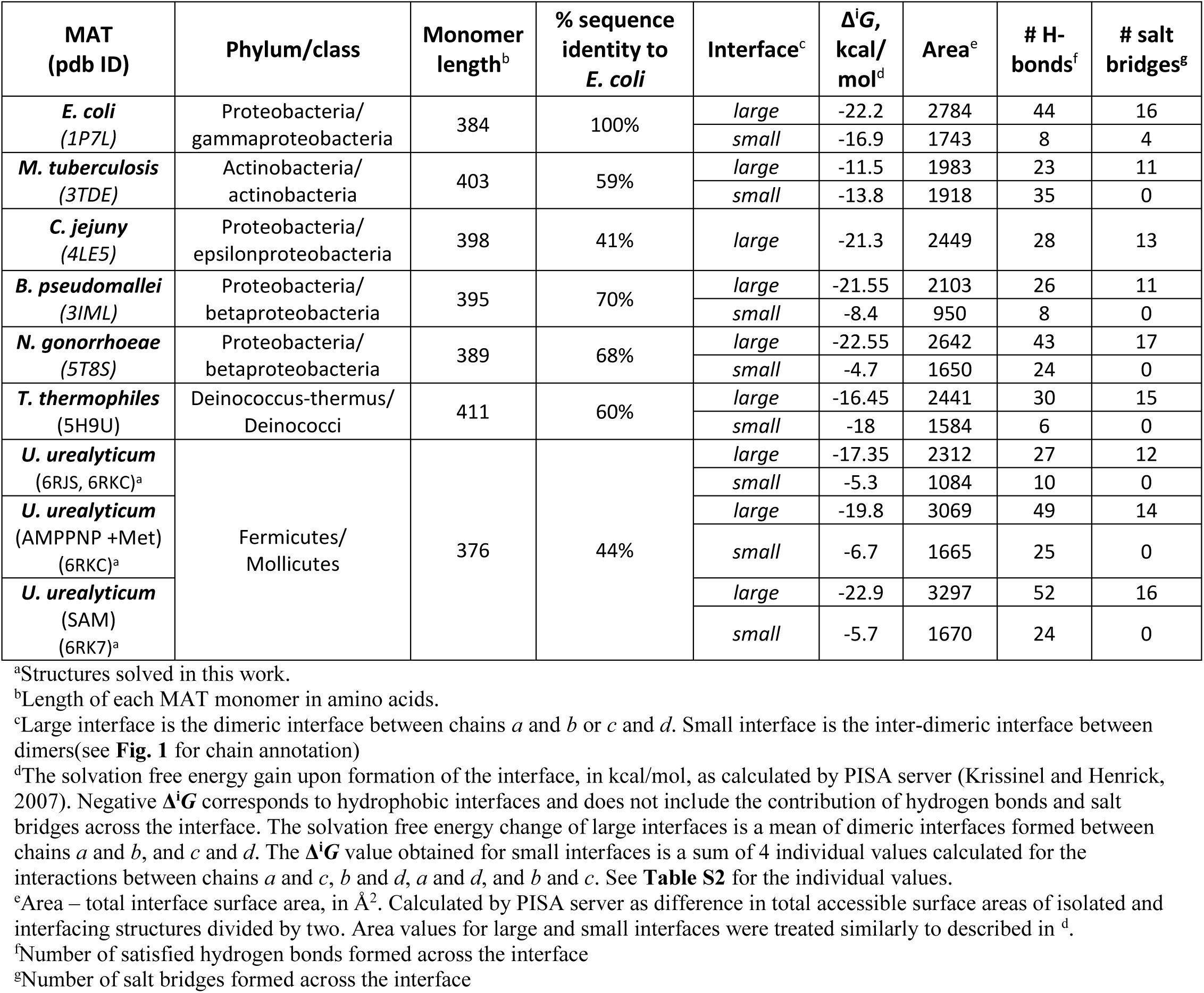
Interface size and composition analysis of orthologous MATs (see also **Table S2**).

**Figure 1.**
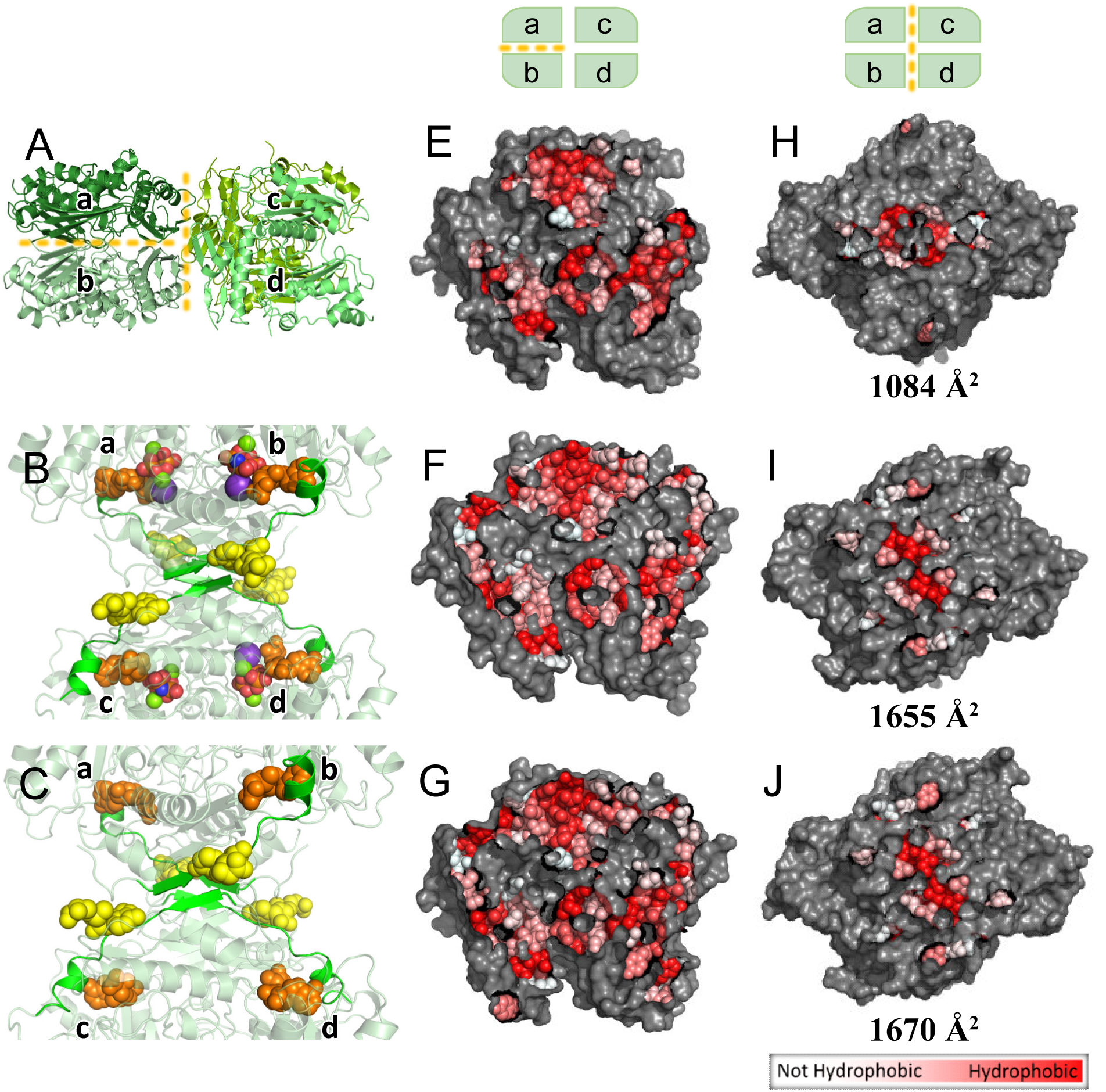
SAM binds to the inter-dimeric interface of uMAT homotetramer. (A) Cartoon representation of uMAT crystal structure in the absence of ligands (PDB ID: 6RJS, 6RKC). See also Figure S1 and Figure S2. uMAT is a dihedral homotetramer formed along two isologous interfaces (shown in yellow dashed lines): Dimeric interface (between chains *a* and *b*, and between chains *c* and *d*), and inter-dimeric interface between dimers *ab* and *cd*. (B) Cartoon representation of uMAT crystal structure formed in the presence of AMPPNP and methionine (PDB ID: 6RKC). Four SAM molecules bind to each of the active sites (orange spheres), and four additional SAM molecules bind to the inter-dimeric interface (yellow spheres). Highlighted in green are the β-strands that participate in formation of continuous β-sheet passing through the inetrface, and flexible loops that adopt a closed state and lock the reaction products, SAM and PPNP, within the active sites. Potassium and magnesium ions are shown as purple and green spheres, respectevly. Phosphate and nitrogen atoms of PPNP are shown as red and blue spheres, respectively (C) Cartoon representation of uMAT crystal structure formed in the presence of SAM (PDB ID: 6RK7). The structure is identical to that shown in (B), with the exception of missing PPNP and metal ions. (E,F,G) Surface representation of the dimeric interface (between chains *a* and *b*) of uMAT structures shown to the left. See also Table 1, and Table S2. Residues directly involved in protein-protein interactions are colored according to their hydrophobicity (Eisenberg et al., 1984). (H,I,J) Surface representation of the inter-dimeric interface (between dimers *ab* and *cd*) of uMAT structures shown to the left. The area of of each interface is designated below (see also Table 1 and Table S2).

A substantially smaller (1,085Å^2^) isologous inter-dimeric interface facilitates the formation of the homotetramer via hydrophobic and H-bonds interactions. In contrary to the large interface, no salt-bridges formed between the dimers (Figure 1H, Table 1, and Table S2). To validate that uMAT is a tetramer in solution, we performed analytical SEC-MALS. At 100 μM (monomer concentration), a major peak of 163 kDA is observed, matching the anticipated molecular weight for uMAT tetramer (Figure S2B). No dissociation is detected with dilution of the protein up to 12.5 μM (3.1 μM tetramer concentration), indicating that uMAT exists in solution predominantly in a tetrameric form at a low micromolar range (Figure S2F).

Analysis of the available structures reveals that almost all bacterial MATs are dihedral homotetramers. Nonetheless, a dimeric MAT from *C. jejuni* (Table 1) is known, suggesting that the homotetramer has evolved by dimerization of the obligatory dimer. Indeed, it was suggested that the evolution of assembly of dihedral homotetramers proceeds via dimeric intermediates and is recapitulated *in vitro* by the (dis)assembly path of the complexes(Levy et al., 2008; Powers and Powers, 2003; Villar et al., 2009). To uncover the (dis)assembly path of uMAT homotetramer, we first subjected the protein to native MS analysis (Figure 2A, **STAR Methods**). Three protein species, tetramer, dimer and monomer, were found, suggesting that uMAT exists at equilibrium between these three states. Next, we subjected uMAT to an equilibrium unfolding in the presence of urea, and monitored the unfolding effects using Trp fluorescence. Over 6M urea range, a three-state unfolding landscape can be observed, with denaturation midpoint (C_m_) of the first transition (between 0 to 3M urea) equals 1.2M, and C_m_ of the second transition (between 3 to 6M urea) equals 4.7M (Figure S2H, **STAR Methods**). Analysis of the oligomeric species of uMAT formed in presence of 0 to 2M urea using SEC-MALS indicated tetramer-dimer dissociation, with approximately half of the species present in either tetrameric or dimeric state around 1M urea (Figure S2A-E). However, a third high molecular weight species was also present, in particular at and above 1.5M urea, suggesting that dimeric and/or monomeric species are prone to aggregation. Collectively, these data indicate that uMAT follows tetramer-dimer-monomer assembly/disassembly path, as anticipated for a dihedral homotetramer.

**Figure 2.**
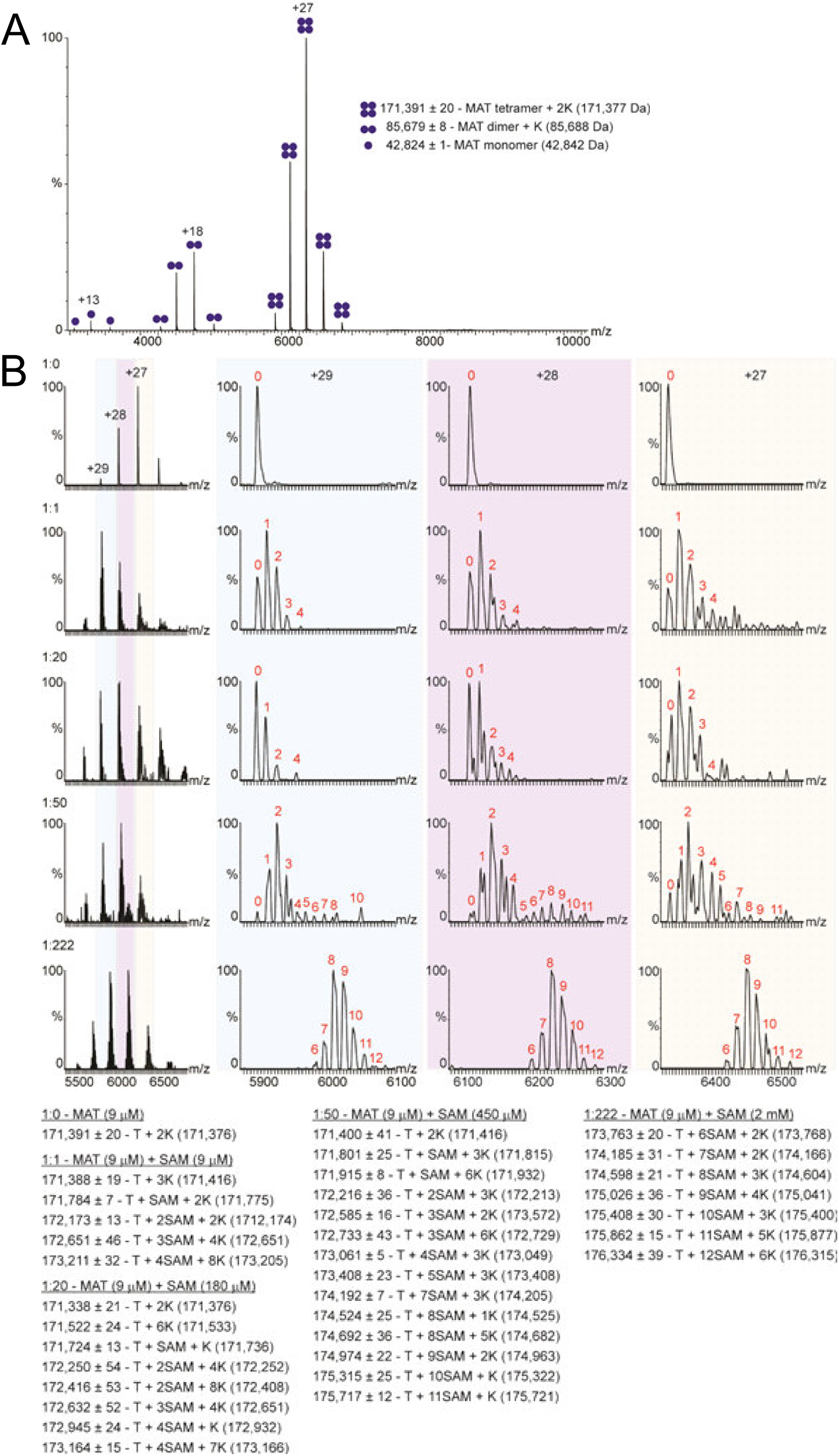
Electrospray native state MS analysis of uMAT. (A) uMAT protein exists in equilibrium between tetramers, dimers and monomers. Mass measurements indicate that the dimer and tetramer are bound to potassium ions (K^+^). Values in brackets show the theoretical masses of the different protein species. (B) The stoichiometry of binding between uMAT and SAM measured by native state MS. uMAT spectra is shown on the left, the enlargement of 3 consecutive charge states (+ 29, +28, +27) are shown on the right. uMAT /SAM molar ratios are indicated on the left side of each tetramer spectrum. The resulting spectra, which are mostly baseline resolved, reveal a clear distribution of peaks corresponding to an ensemble of MAT tetramers bound to an increasing number of SAM molecules and potassium ions. The fine structure of each charge state is a result of the combination of peaks that correspond in mass to different numbers of bound SAM molecules, which are indicated in red. Measured masses of the different species in each experiment are listed under the figure. T – tetramer, K – potassium

### SAM binds into the inter-dimeric interface of uMAT

The reaction catalyzed by MAT proceeds in two, rather unusual steps (Mudd and Cantoni, 1958). First, sulfur of methionine attacks C5’ of ATP, producing SAM and tripolyphosphate (PPP_*i*_). In the second step, the bound tripolyposphate is hydrolyzed into P_*i*_ and PP_*i*_, with P_*i*_ originating from the γ-phosphoryl group of ATP. When ATP is replaced with AMPPNP, the hydrolysis and release of PPP_*i*_ is almost completely blocked. In this case, the dissociation of SAM from the active site is extremely slow (Komoto et al., 2004; Markham et al., 1980). To reveal the structural effects of ligand binding, we solved uMAT crystal structure in presence of AMPPNP and methionine at 2.6Å, and found that all four active sites of the homotetramer are occupied with SAM and PPNP (Figure 1B, Table S1, **STAR Methods**). In addition, the flexible loops (positions 89-116) ─ fully discernable in this structure ─ were found in a closed state, blocking the entrance to the active sites. The structural alignment of uMAT monomer with ordered flexible loop with its counterpart from eMAT reveals that the loop position is almost fully conserved in both structures (Figure S1D). Furthermore, the orientation of SAM within the active sites of uMAT fully overlaps with SAM within the active sites of eMAT (Figure S3A). Accordingly, all contacts between SAM and the active site residues appear fully conserved between MATs with known structures (Figure 3A, Figure S3B,C).

**Figure 3.**
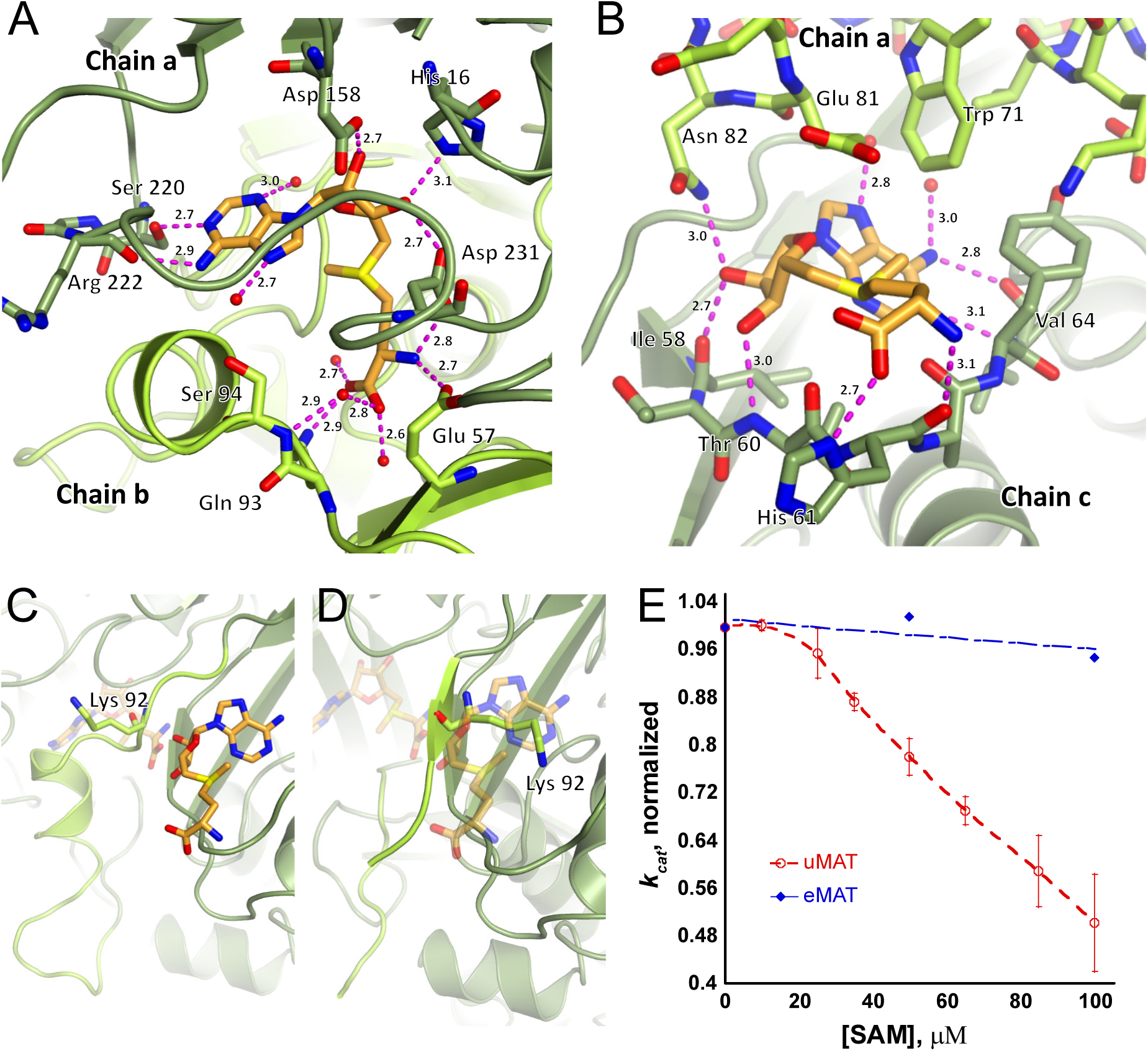
SAM binding to the inter-dimeric interface locks the flexible loop in a closed state. (A) Active site of uMAT tetramer occupied with SAM (PDB ID: 6RK7). Chains *a* and *b* forming the active site pocket are presented as a cartoon. SAM molecule and uMAT residues forming direct interactions with SAM are shown as sticks (nitrogen, oxygen, and sulfur atoms are blue, red, and yellow, respectively). Pink spheres represent water molecules, and dashed pink lines depicts hydrogen bonds. The flexible loop (residues 89-116), shown in light green, adopts a closed conformation and locks SAM molecule within the active site (see also Figure S3). (B) Allosteric binding pocket of SAM in the inter-dimeric interface formed between chains *a* (in light green) and *c* (dark green). SAM (shown in sticks) forms direct interactions with residues from both monomers (also shown in sticks). Water molecules and hydrogen bonds are marked as in (A) (see also Figure S5) (C) SAM binding to the inter-dimeric interface is allowed when the flexible loop (light green) is closed. The faded SAM molecule in the background is occupying the active site. Note the orientation of side chain of Lys 92. (D) The overlay of uMAT structure obtained in the absence of ligands (6RJS) with SAM from the SAM-supplemented structure (6RK7) shows a steric hindrance that prevents SAM binding to the interface when the flexible loop is open. Note the orientation of the side chain of Lys92 that directly clashes with SAM molecule. (E) Interactions between SAM and the inter-dimeric interface pocket. The residues highlighted in light green belong to the opposing dimer. (F) The catalytic turnover (*k_cat_*) of uMAT (shown in red) is highly sensitive to SAM concentration. Under the identical conditions, the catalytic turnover of eMAT (blue) is only slightly affected. Error bars represent standard deviation between three independent measurements. See also Figure S4 and **STAR Methods**.

Although the addition of ligands did not prompt changes in the organization of the dimeric interface (with the exception of the contribution of the resolved active site loops, Figure 1F, Table 1, and Table S2), we found profound changes in the inter-dimeric interface of the homotetramer (Figure 1I, Table 1). First, in addition to the four SAM molecules occupying the active sites, four additional SAM molecules were found bound to the inter-dimeric interface (Figure 1B). Second, dimers have rotated along the interface by ∼45°. This rotation aligned the β-strands of the N-terminal domains into a continuous β-sheet running throughout the interface (Figure 4), and increased the interface surface by almost 60% (from 1,084Å^2^ to 1,665Å^2^) (Figure 1H,I, Table 1, and Table S2).

**Figure 4.**
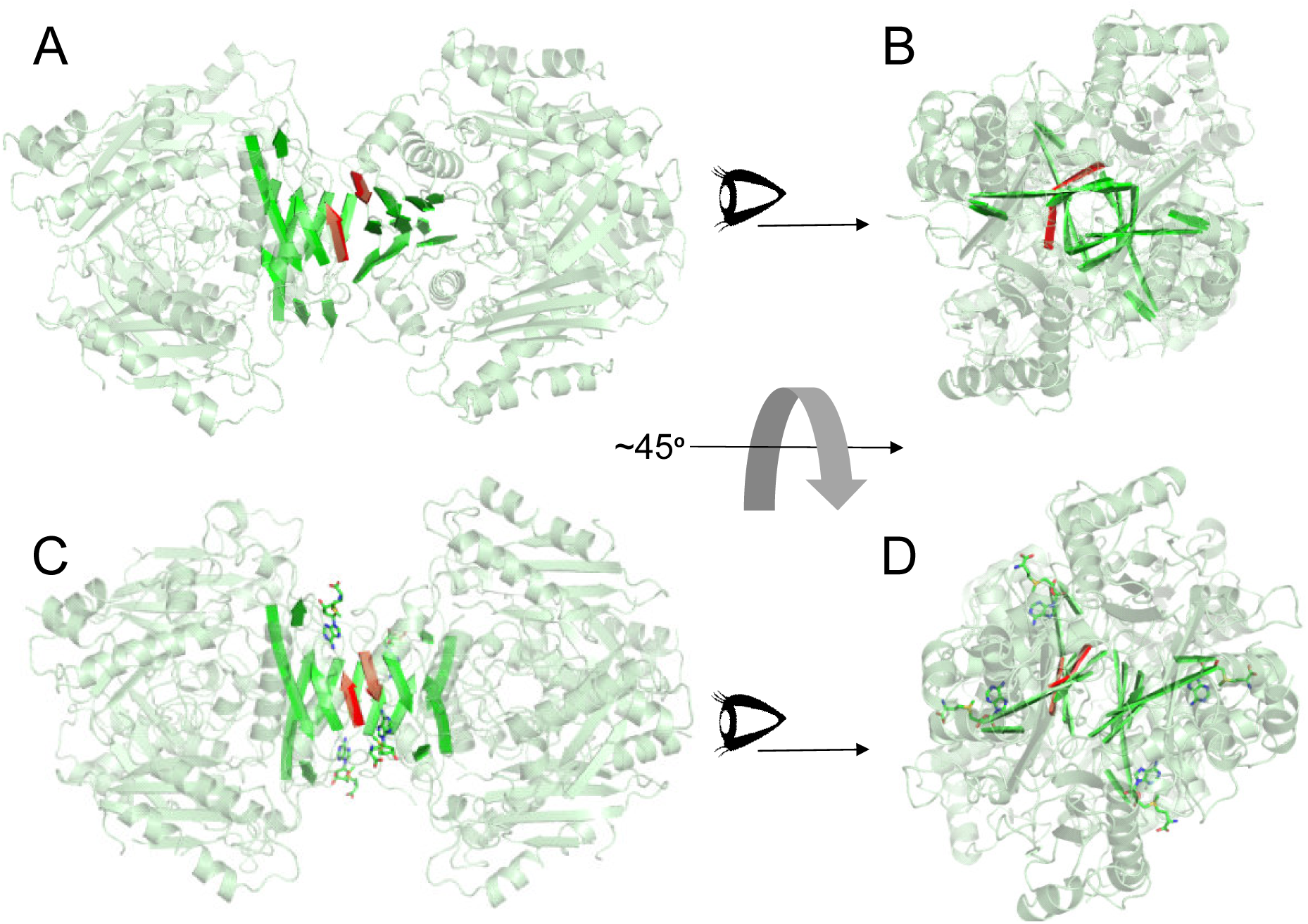
SAM binding to the inter-dimeric interface is accompanied by structural rearrangement of uMAT tetramer. (A) Position of β-strands (colored in red and green) running through the inter-dimeric interface of uMAT obtained in the absence of ligands (PDB ID: 6RK7). (B) Back view of the structure shown in A. (C) When SAM (shown in sticks) is bound to the inter-dimeric interface, dimers rotate by ∼45°C, resulting in formation of a continuous β-sheet that stiches both dimers together (PDB ID: 6RK7). Note the shift in the position of red β-strands. (D) Back view of the structure shown in C.

AMPPNP is not a physiological substrate, and presence of nonhydrolyzable tripoliphosphate in the active site could be partially responsible for the unusual properties of the obtained structure. To uncover the uMAT structural features under physiologically more relevant conditions, we solved uMAT crystal structure in presence of SAM at 1.8Å resolution (Figure 1C**,G,J, Table S1, STAR Methods**). The resulted structure was virtually identical to the structure obtained in the presence of AMPPNP and methionine: four SAM molecules bound to the active sites and sealed behind flexible loops, and four additional SAM molecules bound to the inter-dimeric interface that rotated by ∼45%. (The only obvious difference between the structures is the presence of PPNP in association with two magnesium and one potassium ions in all four active sites, as shown in Figure 1B). Thus, the observed structural features are not a result of nonhydrolyzable PPNP.

### SAM binding to the inter-dimeric interface locks flexible loops and inhibits catalysis

The conformational alternations of the flexible loops gating the access to the active sites (residues 89-116 in uMAT) are necessary to allow access of substrates and departure of products during the catalytic cycle, and are directly linked to the catalytic turnover of MAT, as demonstrated for eMAT (McQueney et al., 2000; Taylor and Markham, 2003; Taylor et al., 2002). Analysis of uMAT structures reveals that binding of SAM molecules to the inter-dimeric interface is sterically possible only when flexible loops adopt a closed conformation (Figure 3B-D). Indeed, in the absence of ligands, the loops are found in open conformations and pass through SAM binding sites in the inter-dimeric interface. The side chain of Lys92, for instance, directly clashes with SAM, thus preventing its binding to the interface when the loop is open (Figure 3C,D). Conversely, the presence of SAM in the active site appears to facilitate loop closure, thus exposing the binding site in the inter-dimeric interface for an additional SAM molecule. This is achieved by a direct hydrogen bond formation between amine moiety of Gln93 side chain of the loop and carboxyl moiety of the active site bound SAM (Figure 3A, Figure S3B). The adjacent Ser94 also forms a hydrogen bond with a water molecule engaged in additional hydrogen bond with the same carboxyl group of SAM (Figure 3A, Figure S3B).

To link this previously unknown mode of interface SAM binding to the catalytic activity of uMAT, we measured uMAT enzymatic activity in the presence and absence of SAM. In the absence of SAM and at a limiting concentration of ATP and saturated amounts of Met, and, conversely, at a limiting concentration of Met and saturated amounts of ATP, we measured the following kinetic parameters: *k_cat_* = ∼0.5 sec^-1^ (per active site), K_m(ATP)_ = 106 μM, and K_m(Met)_ = 51 μM (Figure S4, **STAR Methods**). These values are very close to the kinetic constants reported for eMAT (Markham et al., 1980; Taylor et al., 2002). However, when the enzymatic reaction was conducted at saturated amounts of both ATP and Met (*i.e.*, the reaction proceeded at *V_max_*), addition of SAM to the reaction mix produced a severe reduction in the catalytic turnover (∼50% reduction in *k_cat_* in presence of 100 μM SAM, Figure 3E**, STAR Methods**). In sharp contrast, addition of SAM to eMAT under identical conditions did not produce a significant reduction in *k_cat_* (Figure 3E). This finding is readily explained by the structural rearrangement induced by SAM in uMAT: An increase in SAM concentration leads to locking the active site loops, movements of which are essential for the catalytic turnover.

### SAM binding to the inter-dimeric interface involves interactions with monomers from both dimers

What is the nature of the interactions formed by SAM molecules within the inter-dimeric interface? We found that SAM binding pockets are laid with strong negative charge, which might facilitate the electrostatic interactions with the positively charged sulfonium ion of SAM (Figure S5A). Further, each of the SAM molecules in the inter-dimeric interface forms interactions with monomers from both dimers, thus reinforcing the integrity of the tetrameric complex (Figure 3B). For example, one of the hydroxyl groups of ribose forms hydrogen bonds with the backbone carbonyl of I58 from monomer *c* and side chain carboxyl of N82 from the opposite monomer, *a*. The adenine moiety also interacts with both monomers. Whilst the C-6 amino group of adenine is stabilized by hydrogen bonds formed with backbone of V64 (monomer *c*), W71 from monomer *a* flips to form T-shaped π-stacking interaction with adenine ring (Figure 5A). Accordingly, in solution, addition of SAM is accompanied by a reduction in Trp fluorescence, whereas binding of ATP to the active site is not accompanied by a fluorescence change (Figure 5B). This suggests that the drop in Trp fluorescence upon SAM addition is predominantly originating from the shift in W71 orientation in the interface.

**Figure 5.**
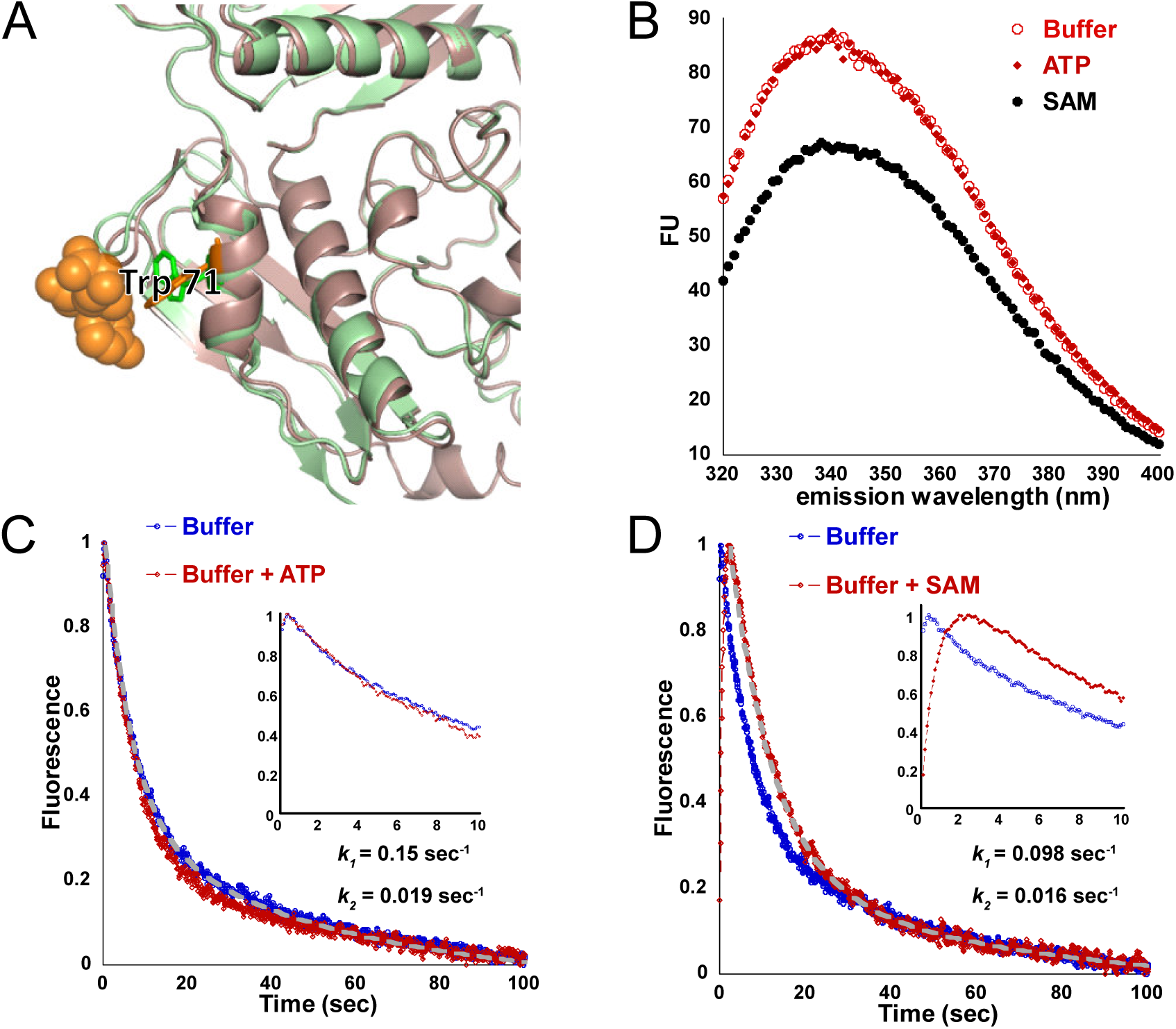
SAM binding to the inter-dimeric interface delays the rate of uMAT tetramer dissociation. (A) Structural alignment between uMAT monomer in the absence (green, PDB ID: 6RK7) and presence (brown, PDB ID: 6RK7) of SAM shows a shift in the orientation of W71. SAM molecule in the interface is depicted as orange spheres. (B) Steady-state tryptophan fluorescence measurement of uMAT in the absence (empty red circles) and presence of ATP (red diamonds) or SAM (black closed circles). Binding of SAM is accompanied by a substantial drop in Trp fluorescence. (C, D) Kinetics of uMAT urea-induced unfolding monitored by a shift in Trp fluorescence in the absence (blue trace) and presence of ATP (red trace, (C)) or SAM (red trace, (D)). The kinetic traces were fitted to double exponential (see **STAR Methods**). Insets show the kinetic traces within the first 10 sec. In the presence of SAM, the kinetics of uMAT tetramer dissociation is markedly delayed.

We could thus utilize Trp fluorescence to measure the effect of SAM binding on the kinetics of uMAT tetrameric dissociation. To this end, we mixed uMAT with 3M urea in a stopped-flow apparatus and monitored the pre-steady-state dissociation kinetics with Trp fluorescence. This particular concentration of urea was chosen, because, as demonstrated above with equilibrium unfolding of uMAT in presence of urea, at 3M urea tetrameric and dimeric species of uMAT are fully dissociated, yet the monomeric species has not yet undergone unfolding (see Figure S2). In the absence of ligands, mixing of uMAT with 3M urea resulted in a rapid drop in Trp fluorescence (Figure 5C). Fitting the trace to a double exponential produced an observed rate constant *k*_obs_ ∼ 0.15sec^-1^ for the first exponential. The value of the observed rate constant for the second exponential was an order of magnitude lower, and, therefore, dismissed (Figure 5C**, STAR Methods**). Addition of ATP did not affect the dissociation kinetics (Figure 5C). However, in presence of SAM, two important changes were observed (Figure 5D). First, a drop in Trp fluorescence was preceded by a rapid trace of ∼2 sec, during which Trp fluorescence has increased. Since association of SAM with the interface is accompanied by a drop in Trp fluorescence (Figure 5B), we interpret the rise in Trp fluorescence to be a result of SAM dissociation from the interface. Importantly, an equimolar amount of ATP did not produce any significant change in the onset of the kinetic trace comparatively to that obtained in the absence of ligands (Figure 5C). Second, fitting the trace of the delayed drop in Trp fluorescence to a double exponential shows ∼30% reduction in the observed rate constant (*k*_obs_ ∼ 0.1sec^-1^) (Figure 5D). These data suggest that binding of SAM molecules to the inter-dimeric interface increases the kinetic stability of uMAT tetramer.

### SAM binding preserves the integrity of the homotetrameric complex

To further explore the effect of SAM binding on stability of uMAT tetramer, we used bis-ANS – a probe for surface hydrophobicity and folding intermediates (Acharya and Rao, 2003; Cardamone and Puri, 1992; Goto and Fink, 1989). Figure 6A shows that addition of both ATP and SAM prompted a substantial reduction in bis-ANS fluorescence at a steady-state, indicating a tighter packing of the protein upon ligand binding. However, the effect of SAM on the reduction of bis-ANS fluorescence was much more pronounced than that of ATP. This difference is expected, since ATP, unlike SAM, is not supposed to bind to the interface, and is therefore unable to trigger the re-alignment of the dimers, and block solvent exposure of the interface.

**Figure 6.**
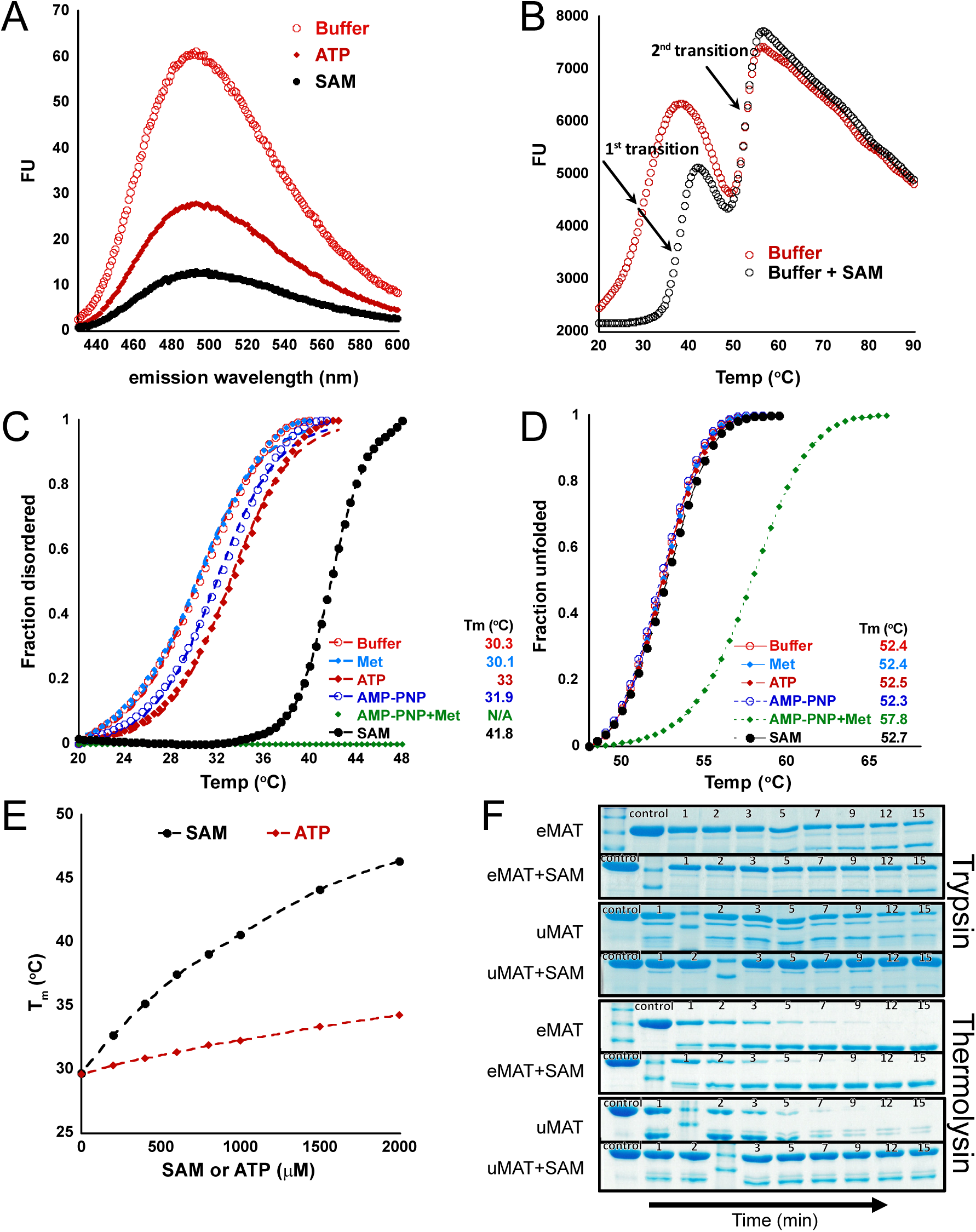
SAM binding stabilizes uMAT tetramer and renders it resistant to proteolysis. (A) Steady state bis-ANS fluorescence measurements of uMAT in the absence (open red circles) and presence of ATP (red diamonds) or SAM (closed black circles). See also Figure S6. (B) SYPRO-orange thermal unfolding of uMAT in the presence (open black circles) and absence (open red circles) of SAM shows that SAM affects the first of the two thermal transitions. (C) Normalized data for the first transition of thermal unfolding shows that SAM (black closed circles) has a great stabilizing effect, whilst a combination of AMPPNP and methionine (green diamonds) fully erases the first transition. To derive the thermal midpoint (T_m_) values, the data were fit to a two-state model (see **STAR Methods**). (D) Normalized data for the second transition of thermal unfolding shows that a combination of AMPPNP and methionine is the only condition affecting the transition. T_m_ values were derived as in C. (E) A correlation between shifts in T_m_ values of the first transition and SAM (black circles) or ATP (red diamonds) concentrations. The effect of SAM on the first transition is profound. (F) Trypsin and thermolysin proteolysis of uMAT and eMAT in the absence and presence of SAM (see **STAR Methods**). Addition of SAM has a profound impact on uMAT resistance to proteolysis, whilst addition of SAM provides no protection from degradation of eMAT.

Next, we used a fluorescence thermal shift assay to follow thermal unfolding of uMAT in a high-throughput manner. To make the assay compatible with a real-time PCR detection system, we replaced bis-ANS with SYPRO-orange dye (see **STAR Methods** and ref. (Chari et al., 2015; Ericsson et al., 2006; Lavinder et al., 2009)). We validated that replacement of bis-ANS with SYPRO-orange indeed produces a pattern of reduction in uMAT fluorescence in presence of SAM or ATP similar to that obtained with bis-ANS (Figure S6A). We followed the thermal unfolding of uMAT at a range of 20°C-90°C (**STAR Methods**). Two distinct transitions can be observed in the absence of ligands (Figure 6B). The thermal midpoint (T_m_) of the second transition (∼50°C) coincides with T_m_ obtained for thermal unfolding monitored by Trp fluorescence, suggesting that the second transition reports the unfolding of monomeric species (Figure 6D and Figure S6B). The much lower T_m_ of the first transition (∼30°C) suggests that it reports on the integrity of the tetramer (order/disorder of the active site loops, and accessibility of the inter-dimeric interface). The addition of AMPPNP and Met has completely erased the first transition (Figure 6C). A similar effect was observed during urea equilibrium unfolding of uMAT in presence of AMPPNP and methionine (Figure S2G,I). This behavior is expected, since production of SAM and PPNP drastically transforms the structural organization of the tetramer: All active site loops are ordered, and the access to the inter-dimer interface is blocked by four SAM molecules. Because SAM and PPNP cannot freely dissociate from the active sites, their presence also shifts the T_m_ of the second transition (monomer unfolding) by ∼5°C (Figure 6D). When added separately, Met and AMPPNP (1 mM each) has no measurable effect on the second transition (Figure 6D). AMPPNP (but not Met), however, has a mild stabilizing effect (Tm increases by ∼2°C) on the first transition (Figure 6C). The effect of ATP addition was almost identical to that of AMPPNP. In contrast, addition of the equimolar amount of SAM had a profound effect on the first transition (Tm increased by >10°C), without having any measurable impact on the second transition (Figure 6C,D). At a range of 0 to 2 mM SAM, the T_m_ of the first transition has shifted by >17°C. The same concentration range of ATP produced only 4°C increase in T_m_ (Figure 6E), indicating that SAM has a profoundly more powerful impact on the integrity of uMAT homotetramer compared to the effect of ATP.

The data presented so far indicate that addition of SAM molecules to uMAT tetramer causes unique structural rearrangement in the tetramer configuration, manifested in highly compact and kinetically stable structure. What is the possible physiological role of SAM-induced kinetic stabilization of uMAT tetramer? It was demonstrated that the kinetic stability of proteins is tightly linked to their ability to withstand proteolysis (Park et al., 2007). We thus compared the sensitivity of uMAT and eMAT to proteolysis by trypsin and thermolysin in the presence and absence of SAM. Whilst no change in eMAT sensitivity to proteolysis was observed, strikingly, addition of SAM practically blocked uMAT degradation by both proteases (Figure 6F). The fact that the structural rearrangement of uMAT tetramer caused by the elevated SAM levels decreases its degradation propensity suggests that SAM levels control uMAT half-life *in vivo*.

### The dynamics of structural rearrangement of uMAT induced by SAM binding is complex

Crystal structures of uMAT, which we solved in the absence and presence of ligands, indicate two distinct homotetrameric states. Multiple structural rearrangements are required to cross between these states, including closure of the flexible loops, re-alignment of dimers, and binding of two SAM molecules per each monomer. Although SAM binding to the interface is sterically blocked when the flexible loop is open, no steric hindrance prevents its binding prior to structural rearrangement of the dimers, further increasing the number of possible structurally distinct conformations existing between these two states (Figure S5B). Collectively, these features suggest that in solution an equilibrium between multiple structurally distinct states should exist at a given concentration of SAM. To test this conjecture, we explored the stoichiometry of SAM binding to uMAT using native state MS (**STAR Methods**). The results show that up to a 20-fold molar excess of SAM, the uMAT tetramer associates with up to 4 molecules of SAM. However, in the presence of increasing SAM concentrations, a step-wise shift of SAM binding to uMAT is observed (Figure 2B). At the highest measured ratio (1:200), the entire population of uMAT shifts towards binding of 6-12 molecules of SAM, suggesting positive cooperativity in SAM binding. (The binding of higher than 8 molecules of SAM to the tetramer is attributed to nonspecific binding, which is a known drawback of the electrospray ionization method (Wang et al., 2003)).

The complex dynamics described above is probably further complicated in the living cell due to competition between SAM and ATP for binding to the active sites. In an attempt to understand the effects of ATP/SAM competition, we solved a crystal structure of uMAT in the presence of both ligands at 2.5Å resolution (Figure S1E, Table S1, and **STAR Methods**). We found that addition of both ligands does not trigger structural rearrangement of uMAT tetramer. The large interface of this structure is identical to the large interfaces of all other uMAT structures (Figure S1F, Table 1, and Table S2), whereas the small interface is identical to that determined for uMAT in the absence of ligands (Figure S1G, Table 1, and Table S2). The active site of chain *a* showed electron-density map of both ATP and SAM, revealing that the obtained crystal structure, in fact, reports an ensemble of states. However, it is noteworthy, that the flexible loop gating the active site (positions 96-108) was resolved only when SAM is present in the active site (chains *a* and *c*, Figure S1E). ATP alone was not sufficient to prompt the closure of the loop (chain *b*, Figure S1E). These features hint to the possibility that there is a kinetic barrier to reach the structural rearrangement of uMAT tetramer: Unless all loops are closed, the dimers cannot realign along the inter-dimeric interface and reach a more stable configuration with markedly increased interface surface area. Reaching this state is possibly the reason for an apparently cooperative SAM binding detected by native state MS (Figure 2B). Since presence of ATP in the active site does not constrain the movement of the flexible loops, it is plausible that when ATP outcompete SAM in the active site, SAM binding to the inter-dimeric interface is hindered. It follows, that ATP/SAM ratio, rather than SAM alone, controls the structural rearrangement of uMAT tetramer *in vivo*.

### The inter-dimeric interface of MATs is an adaptive hotspot

To better understand the possible adaptive role of the inter-dimeric interface in MATs, we analyzed both isologous interfaces of bacterial MATs with available structures (Figure 7, Table 1, Table S2, **STAR Methods**). We find that the residue composition and geometric patterns of the large interface is highly conserved in all MAT structures. This conservation is also manifested in the fact that the surface area of the large interface roughly scales up with the number of H-bonds, which is a general feature of protein interfaces(Janin et al., 1988) (Figure S7A, Table 1, Table S2). Conservation of the large dimeric interface holds despite the fact that bacterial species within which the MAT proteins resign occupy highly distinct ecological niches, including disease-causing parasitizing bacteria (*N. gonorrhoeae* and *U. urealiticum*), and thermophilic bacterium *T. thermophiles*. This finding suggests that the evolution of the large interface is predominantly constrained by the enzymatic activity of uMAT. In sharp contrast, the residue composition, geometric patterns, and the surface area of the inter-dimeric interfaces appear highly diverged (Figure 7, Table 1, Table S2). For example, *E. coli* is the only species among the analyzed MATs, the inter-dimeric interface of which is fortified with salt-bridges. Further, the amount of H-bonds in the inter-dimeric interface does not correlate with the surface area, largely because of the deviation of E. *coli* and *T. Thermophiles* MATs from the general trend (Figure S7B). Overall, these findings support the hypothesis that by acting on the inter-dimeric interface, evolution preserves the enzymatic activity of MATs, while tailoring its other molecular properties, such as the intracellular protein turnover or regulation of activity in response to metabolite levels, to the unique needs of the organism via the tetramer.

**Figure 7.**
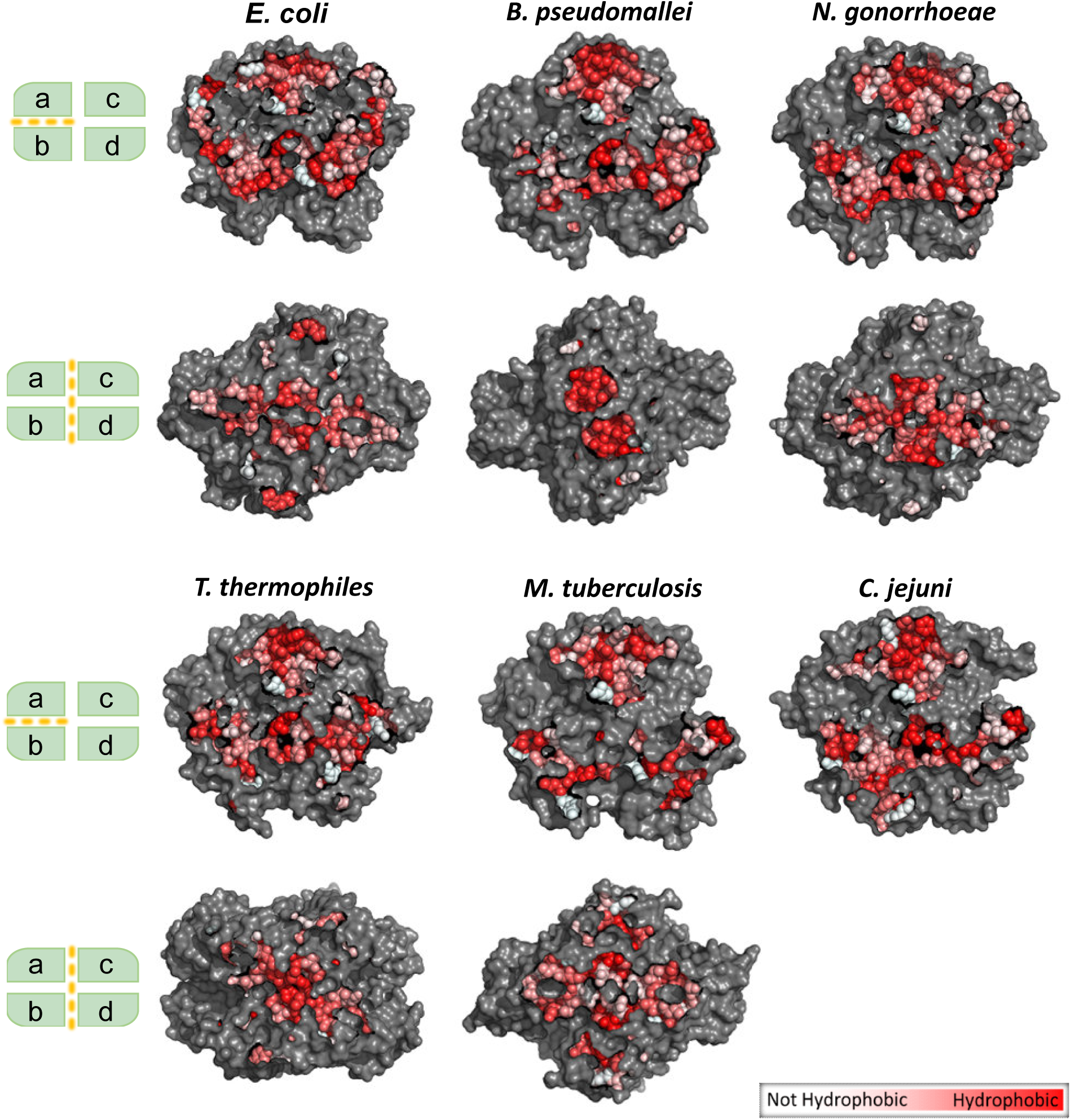
Comparison of large and small interfaces in bacterial MATs with known structures. Surface representation of dimeric (between chains *a* and *b*) and inter-dimeric (between dimers *ab* and *cd*) interfaces. Residues directly involved in protein-protein interactions are colored according to their hydrophobicity(Eisenberg et al., 1984). (See also Table 1 and Table S2). Note the high conservation among large interfaces, and high divergence among small ones. See also Figure S7.

## Discussion

Cellular proteomes are dominated by symmetric homomeric structures (Goodsell and Olson, 2000; Levy et al., 2008; Levy et al., 2006; Marsh et al., 2015), with dihedral homotetramers (dimers of dimers) being the second most abundant symmetry group enriched with metabolic enzymes (Bergendahl and Marsh, 2017; Goodsell and Olson, 2000; Levy et al., 2008). It was suggested that the evolution of assembly of dihedral homotetramers has proceeded via dimeric intermediates (Levy and Teichmann, 2013). However, since the evolutionary more ancient dimeric configurations must already have been functional, the adaptive role of homotetramerization is not obvious, especially, given the fact, that, as is the case with MAT, dihedral homotetramers often have functional dimeric orthologs. Here we address the functionality associated with homotetramerization of MAT from *U. urealiticum*, a dihedral homotetramer, whose crystal structure we solved in the absence and presence of ligands. We show that the evolutionary more recent inter-dimeric interface has evolved to control the enzymatic activity that takes place in the ancient and highly conserved dimeric interface in response to product accumulation. Specifically, the inter-dimeric interface accommodates four allosteric SAM binding sites (one per each monomer). Binding of SAM molecules to the interface is associated with profound structural rearrangements of the entire uMAT tetramer. First, flexible loops gating the access to the active sites remain locked in the closed conformation, thus inhibiting the catalytic turnover. Second, the dimers rotate along the inter-dimeric interface, causing ∼60% increase in the surface area and formation of a continuous β-strand that passes through the interface. Binding of four SAM molecules to the inter-dimeric interface therefore seals the interface from solvent exposure. Together, these structural rearrangements render the entire homotetramer kinetically stable, a property manifested in the markedly decreased sensitivity to proteolysis.

Importantly, neither the allosteric regulation of catalytic activity nor the increased stability of the tetramer in response to SAM accumulation have been reported in other MATs. Mapping the residues directly interacting with SAM in the inter-dimeric interface of uMAT on multiple sequence alignment comprised of bacterial MATs suggests that other Mollicutes, *e.g.*, *M. pneumonia*, might possess a similar regulation mode (Figure 3B and Figure S7C). Why did *U. urealiticum* and, possibly, other Mollicutes evolve this unique mode of regulation? Evolution of Mollicutes was accompanied by massive genome reduction that resulted in reduced metabolic capacity and absence of effectively functioning transcriptional regulation (Maier et al., 2013; Razin et al., 1998; Sirand-Pugnet et al., 2007). For instance, methionine metabolism is tightly regulated by a network of transcription factors and metabolites (including SAM) in *E. coli* (Weissbach and Brot, 1991). This type of regulation is completely missing in *U. urealiticum* that does not possess neither genes involved in *de novo* methionine synthesis nor transcription factors needed for its regulation. The paucity of transcriptional regulation of metabolism in Mollicutes could have resulted in “delegating” the regulation of MAT activity directly to the protein level. In addition, many Mollicutes, including *U. urealiticum*, have lost GroEL/ES chaperonins, but preserved AAA+ ATP-dependent proteases, such as Lon (Clark and Tillier, 2010; Wong and Houry, 2004). This prompted the idea that proteostasis in Mollicutes might be skewed towards protein degradation (Wong and Houry, 2004). Accordingly the increased stability of uMAT tetramer in the presence of saturating amounts of SAM might be an adaptive feature that prevents its degradation *in vivo*.

The regulation of uMAT activity and stability via SAM binding to the inter-dimeric interface of the tetramer is obviously a functionally beneficial result of selection operating on the interface. Analysis of the interfaces of other bacterial MATs with known structures reveals a sharp contrast in the evolutionary preservation of the dimeric versus the inter-dimeric interfaces. Whilst the larger and evolutionary ancient dimeric interface preserves residue composition, types of contacts, and surface geometry across MATs originating from bacteria that occupy distinct ecological niches, the smaller and evolutionary recent inter-dimeric interface exhibit highly diverged surface geometries, residue composition, and nature of contacts within the interface. For instance, the inter-dimeric interface of MAT from *T. thermophiles* has a relatively low number of hydrogen bonds (only 6 per surface of 1584Å^2^), but also the highest number of hydrophobic contacts among all bacterial MATs, manifested in the lowest solvation energy per 1Å^2^ of the inter-dimeric interface (Table 1). Since hydrophobic packing is the major contributing factor to stabilization of thermophilic proteins (Gromiha et al., 2013), highly dense network of hydrophobic interactions in the inter-dimeric interface is likely an adaptation to thermophilic life-style. Based on this analysis, we propose that, whilst the evolutionary ancient and highly conserved dimeric interface of MAT controls its primary enzymatic function and the assembly path of the tetramer, the more recent inter-dimeric interface is recruited by evolution to tune the molecular properties of the entire tetramer and tailor them to the specific needs of the organism. Finally, the existence of the homomer-level modes of regulation of stability and activity that stems from the evolutionary tinkering of the inter-dimeric interface also provides a clue as to why dihedral homomers exhibit a particularly strong functional enrichment in metabolic enzymes (Bergendahl and Marsh, 2017).

## Author contributions

Conceptualization, S.B.; Investigation, D.K., F.S., G.B.-N., A.S.; Formal analysis, R.Z., A.S., G.B.-N., M.S; Visualization, S.B., D.K.; Writing – Original Draft, S.B.; Funding Acquisition, S.B.; Supervision, S.B.

## Acknowledgements

We thank Dan Tawfik and Amir Aharoni for insightful comments and help with preparation of the manuscript. This work was supported by Israel Science Foundation grant 1630/15 to S.B.

## STAR METHODS

### Lead Contact and Materials Availability

Further information and requests for resources and reagents should be directed to and will be fulfilled by the Lead Contact, Shimon Bershtein

### Experimental Model and Subject Details

A bacterial strain used for DNA manipulation was *E. Coli* XL-1 Blue (Stratagene). A bacterial strain used for protein expression was BL21(DE3) (ThermoFisher). Cells were cultured in L-broth supplemented with the appropriate antibiotics.

### Method Details

### Gene cloning, protein expression and purification

The gene encoding MAT from *U. urealiticum* (uMAT) was custom synthesized by was custom synthesized by Integrated DNA Technologies with a fused fragment encoding N-terminal Hisx6 tag and flanking NdeI and XhoI restriction sites. To this end, the protein sequence of uMAT (NCBI reference sequence WP_004025944.1) was converted to DNA sequence using the manufacturer’s *E. coli* codon optimization tool. The gene encoding MAT from *E.coli* (eMAT) was amplified directly from the chromosome of *E. coli* MG1655 using the following primers: For-CATATGGCAAAACACCTTTTTACGTCCGAGTCCGTCTC and Rev-CTCGAGCTATTAATGGTGATGGTGATGGTGCTTCAGACCGGCAGCATCGCGCAGCAGCTGCGCTTTG. The For primer introduces NdeI restriction site. The Rev primer introduces a fused fragment encoding N-terminal Hisx6 and XhoI restriction site. Both genes were cloned into pET24a expression system using NdeI/XhoI restriction sites, and expressed in BL21(DE3) cells. Specifically, an over-night starter was diluted 1:100, grown at 37°C until OD600nm = 0.5, after which the expression was induced by addition of 0.4mM IPTG over-night at 30°C. Cells were centrifuged at 4800xg, and the pellet was stored at -20°C. Cells were lysed by sonication after a 30 min pre-incubation with 1 mg/ml lysozyme (Sigma) and 500U benzonase (Sigma) on ice. The filtered lysate was purified by Ni-NTA on a His-TRAP FF 5ml column (GE Healthcare) and dialyzed into 25mM Tris pH 8.0, 150mM KCl, 1mM DTT. This was followed by size exclusion chromotography on a Superdex 200 16/600 column (GE Healthcare) in the same buffer. uMAT fractions were dialyzed against 20mM MOPS-KOH pH 7.5, 10mM KCl, 1mM DTT, while EcMAT fractions were not subjected to buffer exchange. The proteins were concentrated using Amicon centrifugal filters and stored at 4°C.

### Protein Crystallization, Data Collection, Structure Determination and Refinement

*uMAT structure in the absence of ligands (PDB ID: 6RJS)* 5 mg/ml of uMAT protein was mixed 1:1 (v/v) with reservoir solution, and crystallized by the sitting drop method over a reservoir containing 5% Tacsimate, 0.1M Hepes pH 7.0, and 10% PEG 5K at room temperature. Crystals were harvested, cryoprotected, and flash-cooled in liquid N_2_. X-ray diffraction data were collected at beamline ID30B of the European Synchrotron Radiation Facility (ESRF, Grenoble, France). Data were collected at 100 K from one crystal of uMAT that diffracted to 2.6 Å resolution. The crystal belongs to the space group P1, with unit cell dimensions of a 56.205, b 64.064, and c 114.495, and it contains four copies of the protein in the asymmetric unit. X-ray data were merged and scaled using XDS (Kabsch, 2010) and was solved by molecular replacement using Phaser (McCoy et al., 2007) in CCP4 (Winn et al., 2011). Native monomer of MAT from *E.coli* (Protein Data Bank (PDB) ID: 1FUG) was used as a search model. Refinement employed alternating cycles of manual rebuilding in COOT (Emsley and Cowtan, 2004) and automated refinement using Refmac5 (Murshudov et al., 2011). The coordinates and structure factors have been submitted to the Protein Data Bank under the accession code 6RJS.

*uMAT structure in the absence of ligands (PDB ID: 6RK5).* 10 mg/ml <u>uMAT</u> was mixed 1:1 (v/v) with reservoir solution, and crystallized by the sitting drop method over a reservoir containing 0.2M MgCl_2_, 0.1M Hepes pH 7.4 and 20% PEG 3350 at room temperature. Crystals were harvested, cryoprotected, and flash-cooled in liquid N_2_. X-ray diffraction data were collected at beamline BL14.1 of BESSY II photon source - Helmholtz-Zentrum Berlin (HZB, Berlin, Germany). Data were collected at 100 K from one crystal of uMAT that diffracted to 2.6 Å resolution. The crystal belongs to the space group P21, with unit cell dimensions a 56.358, b 80.450, and c 158.849, and it contains four copies of the protein in the asymmetric unit. X-ray data were merged and scaled using XDS (Kabsch, 2010) and was solved by molecular replacement using Phaser (McCoy et al., 2007) in CCP4 (Winn et al., 2011). Native monomer of MAT from *E.coli* (Protein Data Bank (PDB) ID: 1FUG) was used as a search model. Refinement employed alternating cycles of manual rebuilding in COOT (Emsley and Cowtan, 2004) and automated refinement using Refmac5 (Murshudov et al., 2011). The coordinates and structure factors have been submitted to the Protein Data Bank under the accession code 6RK5.

*uMAT structure in the presence of AMPPNP and methionine (PDB ID: 6RKC)*. 10 mg/ml uMAT with 5mM AMPPNP and 5mM methionine was mixed 1:1 (v/v) with reservoir solution, and crystallized by the sitting drop method over a reservoir containing 0.1M magnesium formate, and 12% PEG 3350, at room temperature. Crystals were harvested, cryoprotected, and flash-cooled in liquid N_2_. X-ray diffraction data were collected at beamline ID23_1 of the European Synchrotron Radiation Facility (ESRF, Grenoble, France). Data were collected at 100 K from one crystal of uMAT that diffracted to 2.56 Å resolution. The crystal belongs to the space group P21, with unit cell dimensions a 142.372, b 79.467, and c 143.787, and it contains eight copies of the protein in the asymmetric unit. X-ray data were merged and scaled using XDS (Kabsch, 2010) and was solved by molecular replacement using Phaser (McCoy et al., 2007) in CCP4 (Winn et al., 2011). Native monomer of MAT from *E.coli* (Protein Data Bank (PDB) ID: 1FUG) was used as a search model. Refinement employed alternating cycles of manual rebuilding in COOT (Emsley and Cowtan, 2004) and automated refinement using Refmac5 (Murshudov et al., 2011). The coordinates and structure factors have been submitted to the Protein Data Bank under the accession code 6RKC.

*uMAT structure in the presence of SAM (PDB ID: 6RK7).* 10 mg/ml uMAT with 1mM SAM was mixed 1:1 (v/v) with reservoir solution, and crystallized by the sitting drop method over a reservoir containing 0.1M Hepes pH 7.0 and 30% Jeffamine ED-2001 at room temperature. Crystals were harvested and flash-cooled in liquid N_2_. X-ray diffraction data were collected at beamline ID29 of the European Synchrotron Radiation Facility (ESRF, Grenoble, France). Data were collected at 100 K from one crystal of uMAT that diffracted to 1.8 Å resolution. The crystal belongs to the space group C21, with unit cell dimensions a 114.00, b 106.14, and c 234.38, and it contained four copies of the protein in the asymmetric unit. X-ray data were merged and scaled using XDS (Kabsch, 2010) and was solved by molecular replacement using Phaser (McCoy et al., 2007) in CCP4 (Winn et al., 2011). Native monomer of MAT from *E.coli* (Protein Data Bank (PDB) ID: 1FUG) was used as a search model. Refinement employed alternating cycles of manual rebuilding in COOT (Emsley and Cowtan, 2004) and automated refinement using Refmac5 (Murshudov et al., 2011). The coordinates and structure factors have been submitted to the Protein Data Bank under the accession code 6RK7.

*uMAT structure in the presence of ATP and SAM (PDB ID: 6RKA).* 10 mg/ml uMAT with 4 mM ATP and 2 mM SAM was mixed 1:1 (v/v) with reservoir solution, and crystallized by the sitting drop method over a reservoir containing 0.2m Li2SO4, 0.1M Tris pH 8.5 and 25% PEG 3350 at room temperature. Crystals were harvested and flash-cooled in liquid N_2_. X-ray diffraction data were collected at beamline ID30B of the European Synchrotron Radiation Facility (ESRF, Grenoble, France). Data were collected at 100 K from one crystal of uMAT that diffracted to 2.5 Å resolution. The crystal belongs to the space group I41, with unit cell dimensions a 114.47, b 114.47, and c 228.12, and it contained four copies of the protein in the asymmetric unit. X-ray data were merged and scaled using XDS (Kabsch, 2010) and was solved by molecular replacement using Phaser (McCoy et al., 2007) in CCP4 (Winn et al., 2011). Native monomer of MAT from *E.coli* (Protein Data Bank (PDB) ID: 1FUG) was used as a search model. Refinement employed alternating cycles of manual rebuilding in COOT (Emsley and Cowtan, 2004) and automated refinement using Refmac5 (Murshudov et al., 2011). The coordinates and structure factors have been submitted to the Protein Data Bank under the accession code 6RKA.

### Enzymatic activity

uMAT activity was determined at 37°C in activity buffer (25mM Hepes pH 7.5, 100mM KCl, 10mM MgCl_2_, 1mM DTT) at either a saturated concentration of methionine (5 mM) and a range of ATP concentrations (0 to 500 μM), or at a saturated amount of ATP (5 mM) and a range of methionine concentrations (0 to 500 μM). uMAT (100nM) was pre-incubated with 5mM of either ATP or methionine at room temperature for 30 minutes. The enzymatic reaction was initiated by adding either ATP or methionine. Aliquots were removed from the reaction mix at various time points, and the reaction was stopped by mixing with 10% perchloric acid in a 1:1 ratio. The aliquots were then centrifuged, and the supernatant was separated by HPLC using the MultoHigh SCX 5μ 250×4.6mm column (CS-Chromatographie Service GmbH). The mobile phase consisted of 400mM ammonium fumarate (adjusted to pH 4.0 using formic acid) at a flow rate of 1ml/min, while measuring absorbance at 254nm. Data analysis was performed by integrating the peaks corresponding to SAM, and fitting them to a calibration curve (see Figure S9A,B). The kinetic constants were derived by fitting the resulted data point to a Michaelis-Menten equation (V_max_·[S]/K_m_+[S]) (Figure S9C,D).

To measure the inhibitory effect of the reaction product on uMAT and eMAT, enzymes (100 nM) were pre-incubated with 5 mM methionine and a range of SAM concentrations (0 to 100 μM). The reaction was initiated by adding 1 mM ATP. Since at these highly saturated concentrations of substrates the reactions catalyzed by both eMAT and uMAT proceed at maximal speed, we treated the linear slopes of the initial velocities obtained at each concentration of SAM as *V_max_*. The catalytic turnover numbers (*k_cat_*) were derived by dividing the maximal velocity values by total enzyme concentrations (Figure 2E).

### Analytical SEC-MALS

Multi-angle light scattering (MALS) analysis on MiniDawn TREOS + OPTILAB T-reX (WYATT) was used to determine the molecular weight of uMAT species separated by size-exclusion chromatography (SAM) on AKTA Pure M25 (GE). uMAT samples were incubated for 24h at room temperature in 20mM MOPS-KOH pH 7.5, 10mM KCl, 1mM DTT at different protein concentrations (12.5-200 µM, monomer concentration), or at 100µM in the presence of a range of urea concentrations (0-6M). The samples were then injected into a Superdex 200 Increase 30×1cm column equilibrated with 25mM Tris-HCl pH 8.0, 150mM KCl, 1mM DTT and the corresponding urea concentration.

### Urea equilibrium unfolding experiments

uMAT samples (2µM each) were prepared in 20mM MOPS-KOH pH 7.5, 10mM KCl, 1mM DTT, 10 mM MgCl_2_ at a range of urea concentrations (0-6M) in the absence of ligands, or in the presence of either 1 mM SAM, or 1mM of each AMPPNP and methionine. The samples were incubated for 24 hours at room temperature, and their tryptophan fluorescence (ex. 295nm, em. spectrum 320-400nm) was obtained using Cary Eclipse fluorescent spectrophotometer, Agilent. To account for the shifts in fluorescence, the spectra were analyzed as described in (Rietveld and Ferreira, 1998). Briefly, fluorescence spectral centers of mass at each concentrations of urea were calculated as λ_[Urea]_ = ∑λI(λ)/∑I(λ), where λ is the emission wavelength, and I(λ) is the fluorescent intensity at wavelength λ. The data were globally fitted to apparent two-state (for AMPPNP+methionine) or three-state (for no ligands, or in the presence of SAM) models of protein unfolding, as described in(Walters et al., 2009). Briefly, the two-state model fit was performed using the following equation:

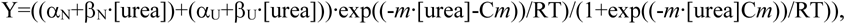

where Y is the apparent fraction of the unfolded protein, α_N_ and β_N_ are the native state signal at zero denaturant, and the slope of the native state baseline, αU and βU are the denatured state signal at 6M urea, and the slope of the denatured state baseline, *m* is the transition slope (dependence of the free energy change on urea concentration, cal/mol/M), *C_m_* is the concentration of urea at the midpoint transition, R is the ideal gas constant (1.987 cal/mol/K), and T is 298K.

The three-state model fit was performed using the following equation:

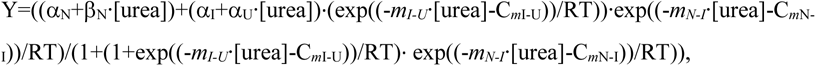

where α_I_ accounts for the signal change at the pre-intermediate state, *m_I-U_* and *m_N-I_* are the slopes of transition between intermediate and denatured states, and between native and intermediate states, and C_*m*I-U_ and C_*m*N-I_ are urea concentrations at midpoint transition between intermediate and denatured states, and between native and intermediate states.

### uMAT urea-induced dissociation kinetics

uMAT samples (4µM) were prepared in Hepes pH 7.5, 10mM KCl, 1mM DTT, and 10mM MgCl_2_ without ligands, or in the present of either 0.4mM ATP or 0.4mM SAM. The uMAT dissociation kinetics was measured in SX20 stopped-flow (Applied Photosystems) by mixing the protein in 1:1 ratio with 6M urea (3M urea final concentration) prepared in the same buffer with a corresponding additive. The ensuing perturbation in tryptophan fluorescence (ex. 295 nm, em. 343 nm) was measured, and the obtained signal was fitted to a sum of two exponentials, as described (Walters et al., 2009):

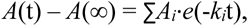

Where *A*(t) is the amplitude at time t, *A*(∞) is the offset value, *A_i_* is the change in signal for phase *i*, and *k_i_* is the observed rate at phase *i*.

### Bis-ANS and Sypro-Orange fluorescence measurements

The effects of ligands on the surface hydrophobicity and folding state of uMAT was probed with 4′-Dianilino-1,1′-binaphthyl-5,5′-disulfonic acid (bis-ANS) (Sigma) and SYPRO-Orange (Sigma) fluorescence dyes. 2µM UuMAT in 20mM MOPS-KOH pH 7.5, 10mM KCl, 10mM MgCl_2_, 1mM DTT without ligands, or pre-incubated for 1h st room temperature with either 5mM ATP or 1 mM SAM were mixed with 10µM bis-ANS or x20 SYPRO-orange dyes. Fluorescence emission spectra were acquired with Cary Eclipse fluorescence spectrophotometer (Agilent) (bis-ANS: ex. 395nm, em. 540-650nm; SYPRO-orange: ex. 490nm, em. 440-600 nm).

### ThermoFluor assay

2µM uMAT samples were prepared in 20mM MOPS-KOH pH 7.5, 10mM KCl, 10mM MgCl_2_, 1mM DTT and x20 SYPRO-orange dye in the presence of one of the following: 1mM ATP, 1mM Methionine, 1mM AMPPNP, 1mM AMPPNP + 1mM methionine, 1mM SAM, or no additives. All samples were pre-incubated for 30 minutes at room temp. and transferred to hardshell PCR plates (BioRad). Fluorescence spectra were obtained in CFX96 Touch™ Real-Time PCR machine (BioRad) using HEX channel (ex. 515-535 em. 560-580) with a 1°/min temperature ramp. The data were analyzed according to described in (Lavinder et al., 2009). Two thermal transitions, 20-45°C, and 50-65°C, corresponding to disintegration of the tetramer, and unfolding of the monomer, respectively, were treated separately. The obtained spectra were normalized and globally fitted to:

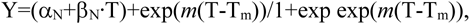

where, Y is the normalized signal, α_N_ and β_N_ are the intercept and slope of baseline for the folded state, and *m* is exponential factor related to the slope of the transition at the apparent melting temperature, *T_M_*.

### Thermal denaturation of uMAT

2uM uMAT in 20mM MOPS-KOH pH 7.5, 10mM KCl, 10mM MgCl_2_, 1mM DTT were melted by 1°C/min temperature ramp in Cary Eclipse fluorescent spectrophotomer (Agilent), and the ensuing change in tryprophan fluorescence was followed (ex. 293 nm, em. 340 nm). The obtained signal was globally fitted to a two-state thermal unfolding model as described in (Jackson and Fersht, 1991):

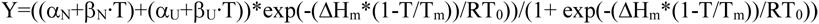

where, Y is the normalized signal, αNand βN are the intercept and slope of baseline for the folded state, αU and βU are the intercept and slope of baseline for the denatured state, Δ Hm is the enthalpy of denaturation at the transition midpoint, T_m_ is the midpoint of thermal denaturation. R is the ideal gas constant (1.987 cal/mol/K), and T0 is 298K. The change in the heat capacity was assumed to be zero.

### MSA

The multiple sequence alignments were produced using the T-COFFEE (Expresso) server(Notredame et al., 2000), and visualized using BioEdit v7.0.5.

### Structural alignment

Structural alignments were performed with PyMol. The alignment of uMAT and eMAT monomers covered 240 amino acids, out of 378 residues for uMAT monomer and 383 residues for eMAT monomer, and resulted in rmsd = 1.35Å. For comparison, alignment of uMAT monomers from the same structure that covered 352 residues out of 378, resulted in rmsd = 0.73Å. Structural alignment of uMAT dimer (PDB ID: 6RJS, 6RK5) and eMAT dimer (PDB ID: 1P7L) produced rmsd = 3.5Å. The alignment covered 280 amino acids. For comparison, alignment of uMAT dimers from the same structure that covered 712 residues resulted in rmsd = 0.99Å.

### Interface analysis (PISA)

Interfaces of various MATs were characterized using protein interfaces, surfaces, and assembly (PISA) server (Krissinel and Henrick, 2007). The interface area is calculated by PISA as a difference in total accessible surface areas of isolated and interfacing structures divided by two. Change in the solvation free energy upon formation of the interface, **Δ^i^G** in kcal/mol, is calculated as a difference in total solvation energies of isolated and interfacing structures. Negative **Δ^i^G** corresponds to hydrophobic interfaces and does not include the effect of satisfied hydrogen bonds and salt bridges across the interface. PISA also predicts hydrogen-bonds and salt-bridges formation in the interface. The inter-dimeric interface of uMAT is comprised of four interacting surface patches. All the parameters pertaining to the inter-dimeric interface are presented as summation over four individual patches (Table 1). The individual parameters of each patch are summarized in Table S2.

### MAT proteolysis assay

uMAT and eMAT proteins were incubated for 30 minutes at room temperature, in the presence or absence of SAM, prior to the addition of either Trypsin or Thermolysin. Final reaction conditions were: 0.02mg/ml Trypsin / Thermolysin, 20mM Tris pH 8.0, 10mM CaCl_2_, 50mM NaCl, 0.5mg/ml MAT, 0.04mM SAM (if added). The reaction was conducted at 37°C and terminated at various time points by boiling the reaction aliquots in SDS loading buffer (40 mM Tris pH6.8, 1mM SDS, 5% glycerol, and 50 mM DTT). Samples were run on 12.5% SDS-PAGE gel, and stained using Coomassie Instant Blue (Expedeon).

### Native state MS

Methionine S-adenosyltransferase (MAT) from *U. urealiticum* was diluted to 18 µM (monomer concentration) in 150 mM ammonium acetate at pH 8 (titrated with ammonia), and buffer exchanged into the same solution. S-adenosylmethionine (SAM, 20 mM, dissolved in 10mM sulfuric acid) was diluted according to need in 150 mM ammonium acetate pH 8. For supplementation of the enzyme with SAM, 2 µl protein were mixed with 2 µl of different concentrations of SAM, in order to reach the required fold excess of MAT:SAM, and analyzed immediately.

Mass spectrometry (MS) measurements were done on a modified Q Exactive Plus EMR Orbitrap (Thermo) (Ben-Nissan et al., 2017). Samples were loaded into gold-coated nano-ESI capillaries, prepared in-house, as previously described (Kirshenbaum et al., 2010) and sprayed into the mass spectrometer. Instrumental parameters included capillary voltage 1.7 kV, inlet capillary temperature of 180 °C, fore-vacuum pressure of 1.42 mbar. Trapping gas pressure was set to 2, corresponding to HV pressure of 5.55×10^-5^ mbar and UHV pressure of 1.48×10^-10^ mbar. Bent flatapole DC bias and gradient were set to 2.2 V and 25 V, respectively, and the HCD cell was operated at 40 V. Mass assignments were done using the SUMMIT 1.0 software (Taverner et al., 2008). Searches were restricted to complexes containing up to four MAT monomers, up to 12 SAM molecules and up to 8 potassium ions, with an error of up to 50 Da.

The stoichiometry of binding between uMAT and SAM was measured under constant uMAT concentration (9 µM monomer concentration) with SAM supplemented at increasing concentrations ranging from 0 to 2mM. The binding of higher than 8 molecules of SAM to the tetramer is attributed to nonspecific binding, which is a drawback of the electrospray ionization method(Wang et al., 2003).

### Quantification and Statistical Analysis

The pre-steady-state kinetics measurements were performed 3-6 times per each condition and were found highly reproducible. The ThermoFluor assay was performed with 3-6 replicates per condition and found to be highly reproducible. Data obtained for uMAT enzyme activity in the presence of SAM represent the mean of three independent measurements (+/-standard error)

### Data and Code Availability

The atomic coordinates of uMAT in the absence of ligands have been deposited in the Protein DataBank Under ID codes 6RJS, 6RK5. The atomic coordinates of uMAT in the presence of AMPPNP+Met, SAM, and ATP+SAM were deposited under ID codes 6RKC, 6RK7, 6RKA.

---

**Figure S1. Related to Figure 1.**
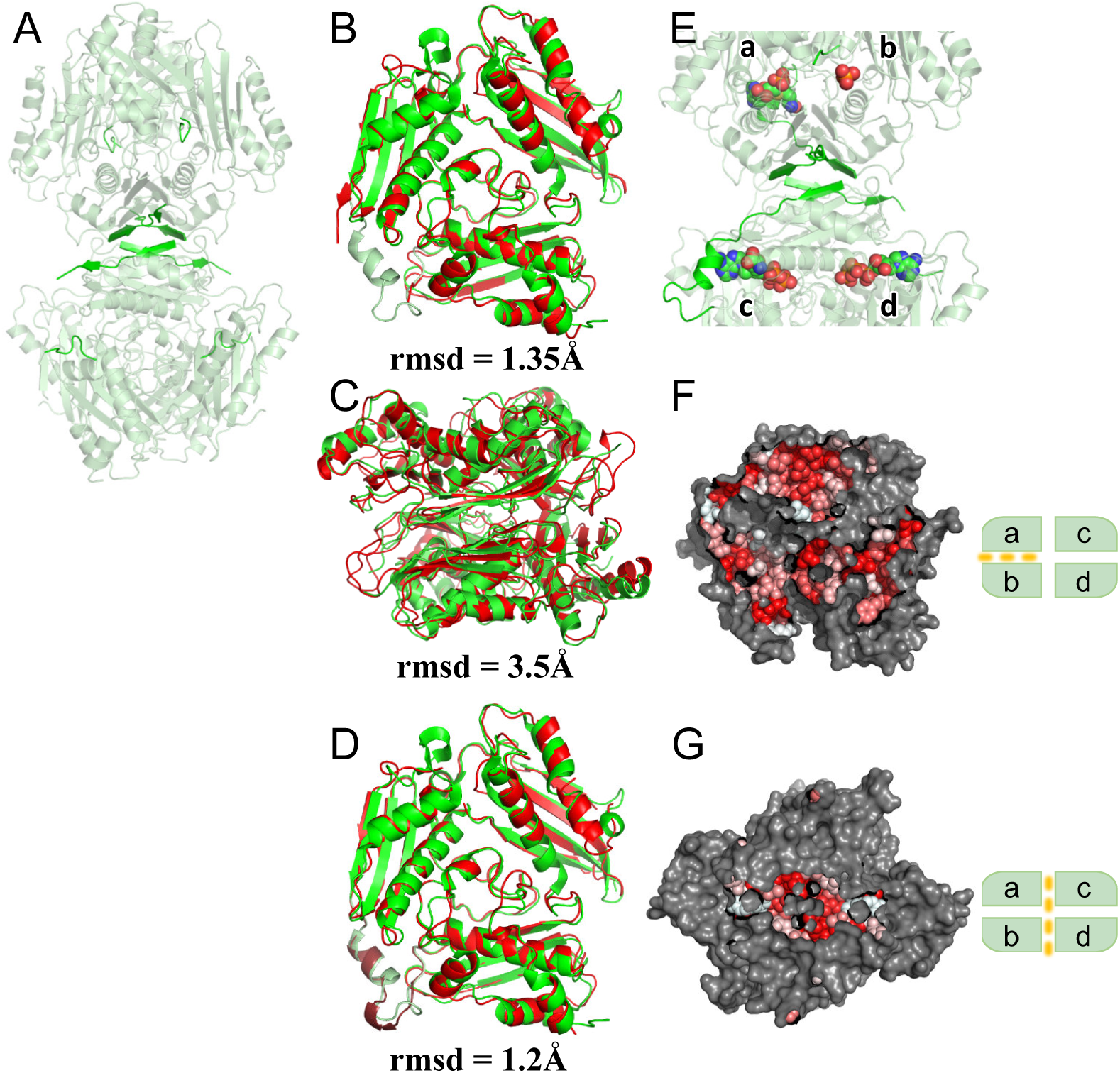
(A) uMAT structure in the absence of ligands (PDB ID: 6RJS, 6RK5). Highlighed in green are the N-terminal β-strands directly participating in formation of the inter-dimeric interface, and both ends of the unresolved flexible loops gating the access to the active sites. (B) Structural alignment of uMAT monomer (red) solved in the absence of ligands (PDB ID: 6RJS, 6RK5) and eMAT monomer (red) solved in the presence of AMPPNP and methionine (PDB ID: 1P7L). Note the missing flexible loop in the uMAT monomer. (C) Structural alignment of uMAT dimer (red) (PDB ID: 6RJS, 6RK5) and eMAT dimer (green) (PDB ID: 1P7L) produced rmsd = 3.5Å. (D) Structural alignment of uMAT monomer (red) solved in the presense of SAM (PDB ID: 6RK7) and eMAT monomer (red) solved in the presence of AMPPNP and methionine (PDB ID: 1P7L) resulted in rmsd = 1.2Å. Note that the fully resolved flexible loop in the uMAT monomer (pale green) largely overlays with the flexible loop of eMAT. (E) cartoon representation of uMAT tetramer. Highlighed in green are the N-terminal β-strands directly participating in the formation of the inter-dimeric interface, and the adjacent flexible loops gating the access to the active sites. Carbon, nitrogen, oxygen, sulfur, and phosphate atoms of the ligands are shown in green, blue, red, yellow, and orange spheres, respectively. Both ATP and SAM molecules can be detected in the active site of chain *a*, suggesting that the obtained structure is an assemble of different conformation. Active site of chain *c* is occupied with SAM and inorganic phosphate. Active sites of chains *b* and *d* harbors ATP and inorganic phosphate molecules, respectively. Note that flexible loops are fully resolved only when SAM is present in the active site (chaind *a* and *c*). (F) Surface representation of the dimeric interface (between chains *a* and *b*). C. Surface representation of the inter-dimeric interface (between dimers *ab* and *cd*). See also Table 1,and Table S2. Residues directly involved in protein-protein interactions are colored according to their hydrophobicity.

**Figure S2. Related to Figure 1.**
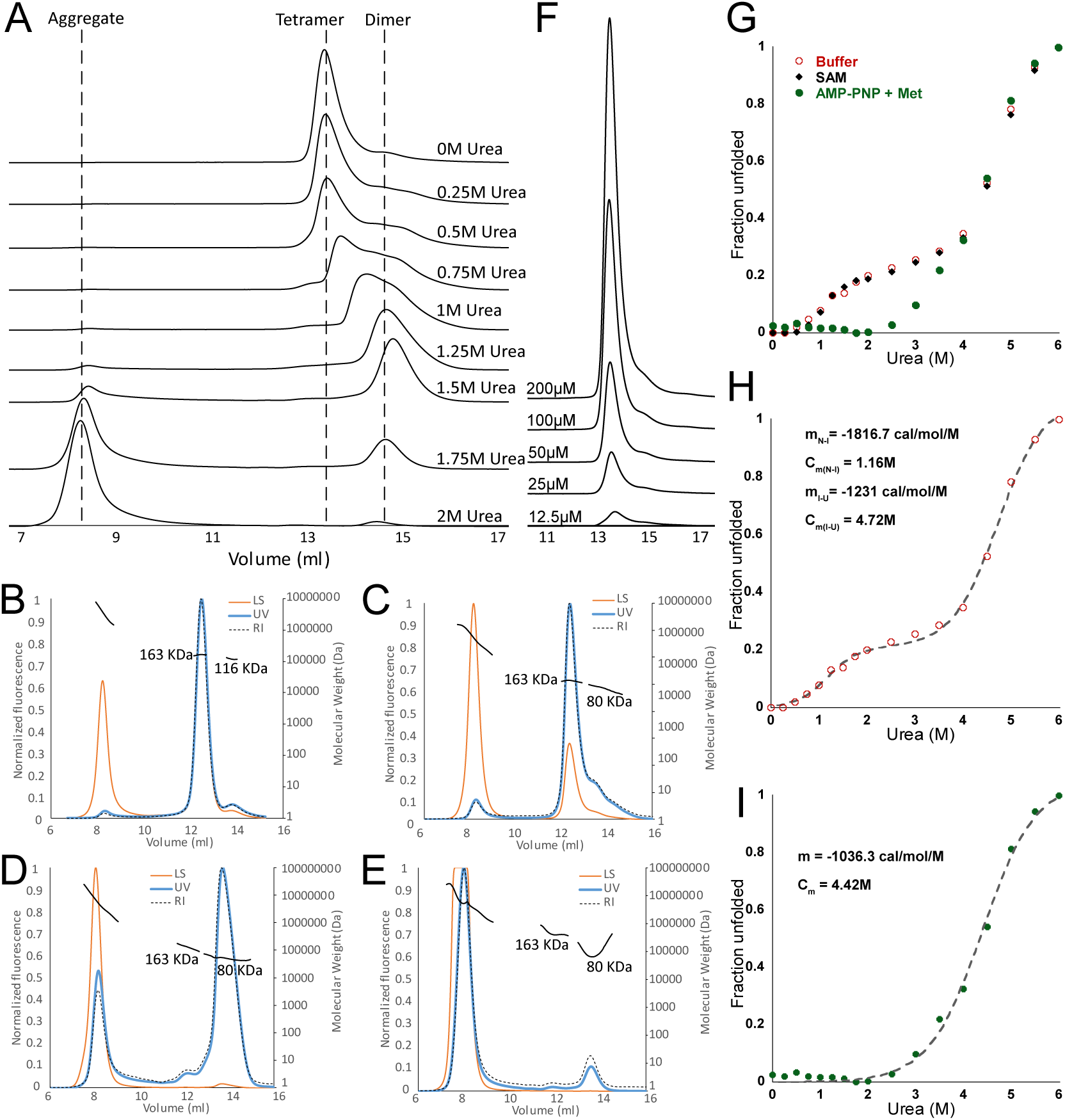
(A) uMAT was pre-incubated at a designated concentration of urea and the obtained oligomeric species were analyzed by SEC (see **STAR Methods**). Between 0 to 1.5Murea, uMAT is found predominantly in either tetrameric or dimeric states. At and above 1.5M urea, and high molecular weight species appear, indicating that the dimeric species tends to aggregate. The molecular weight of the oligomeric species was obtained using SEC-MALS analysis at 0 (B), 0.5 (C). 1.5 (D), and 2 (E) M urea. The static light scattering trace (LC) is in yellow, the UV trace at 280 nm is at blue, and refractive index (RI) is at dashed black (see **STAR Methods**). The weight of the tetrameric uMAT species is ∼163 kDA, the dimeric weight is ∼80 kDa. The 116 kDA peak at 0 urea corresponds to impurity. (F) uMAT was diluted up to 12.5 uM (monomer concentration), pre-incubated for 18 hours, and analyzed on SEC. See **STAR Methods**. Only tetrameric species can be detected. (G) uMAT urea-induced equilibrium unfolding in the absence (open red circles) and presence of SAM (black diamonds), or a combination of AMPPNP and methionine (closed green circles). Note that the unfolding signals in the absence of ligands or in the presence of SAM fully overlaps. The data are normalized as a fraction of unfolded protein. See also **STAR Methods**. (H) Fitting of the uMAT unfolding data in the absence of ligands shown in (G) to a three-state protein unfolding model (see **STAR Methods**). (I) Fitting of the uMAT unfolding data of uMAT in the presence of AMPPNP and methionine shown in (G) to a two-state unfolding protein model (see **STAR Methods**).

**Figure S3. Related to Figure 3.**
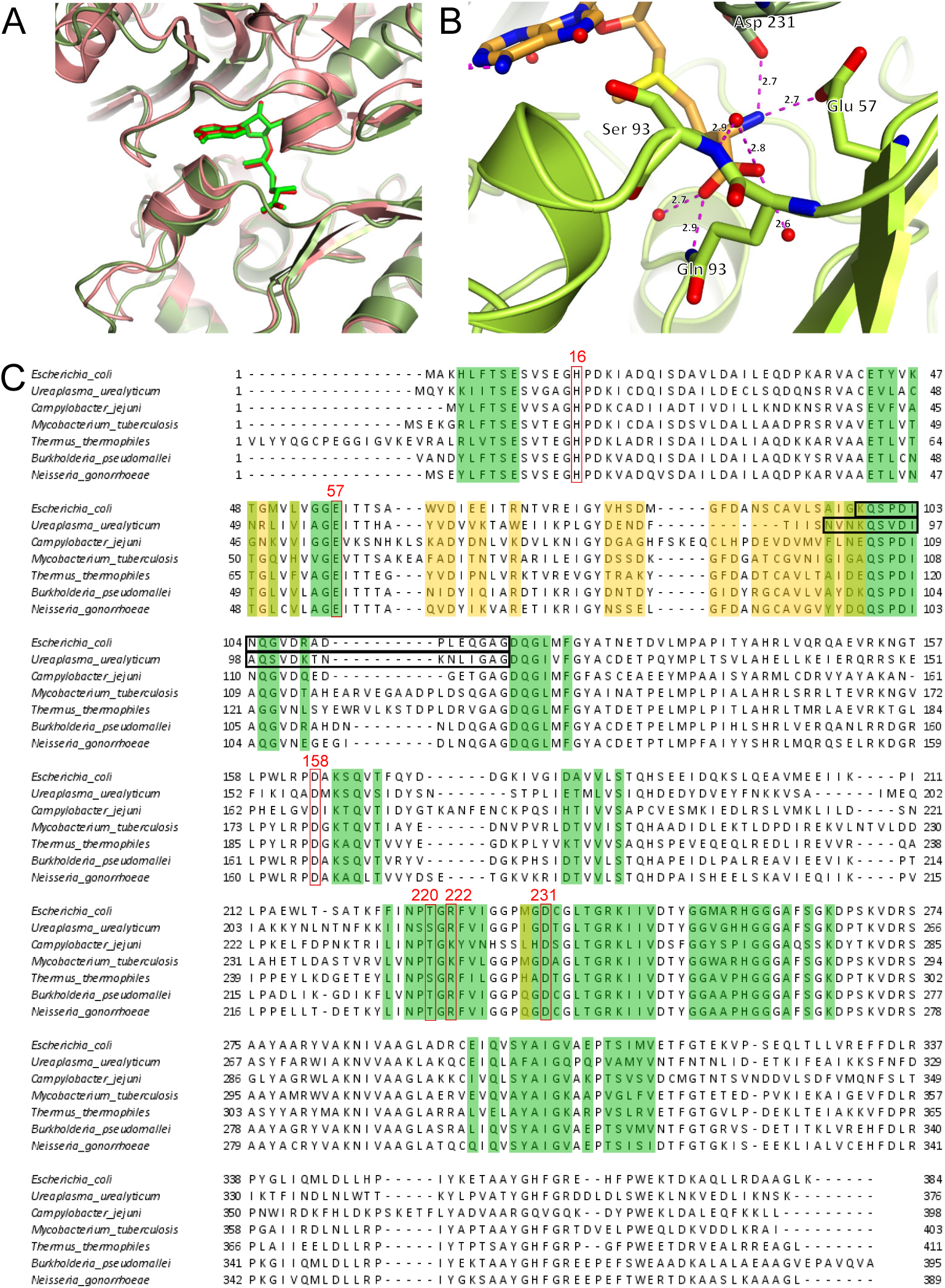
(A) Structural alignment of uMAT (6RK7, pink) and eMAT (P7L, light green) with their respective SAM molecules (shown in sticks) in the active sites. (B) Flexible loop directly interacts with SAM in the active site of uMAT (PDB ID: 6RK7). The flexible loop is shown in light green. SAM and residues interacting with SAM are shown as sticks. Hydrogen bonds are indicated in dashed pink lines. Water molecules are shown as red spheres. (C) Multiple sequence alignment of bacterial MATs. Highlighted in green are residues forming the large dimeric interface in uMAT. Residues that form direct contacts with SAM in the interface are outlined in red and numbered according to uMAT sequence. Highlighted in yellow are residues contributing to the inter-dimeric interface of uMAT. Note the high conservation of the residues contributing to the large interface, and high divergence of the residues forming the small inter-dimeric interface.

**Figure S4. Related to Figure 3.**
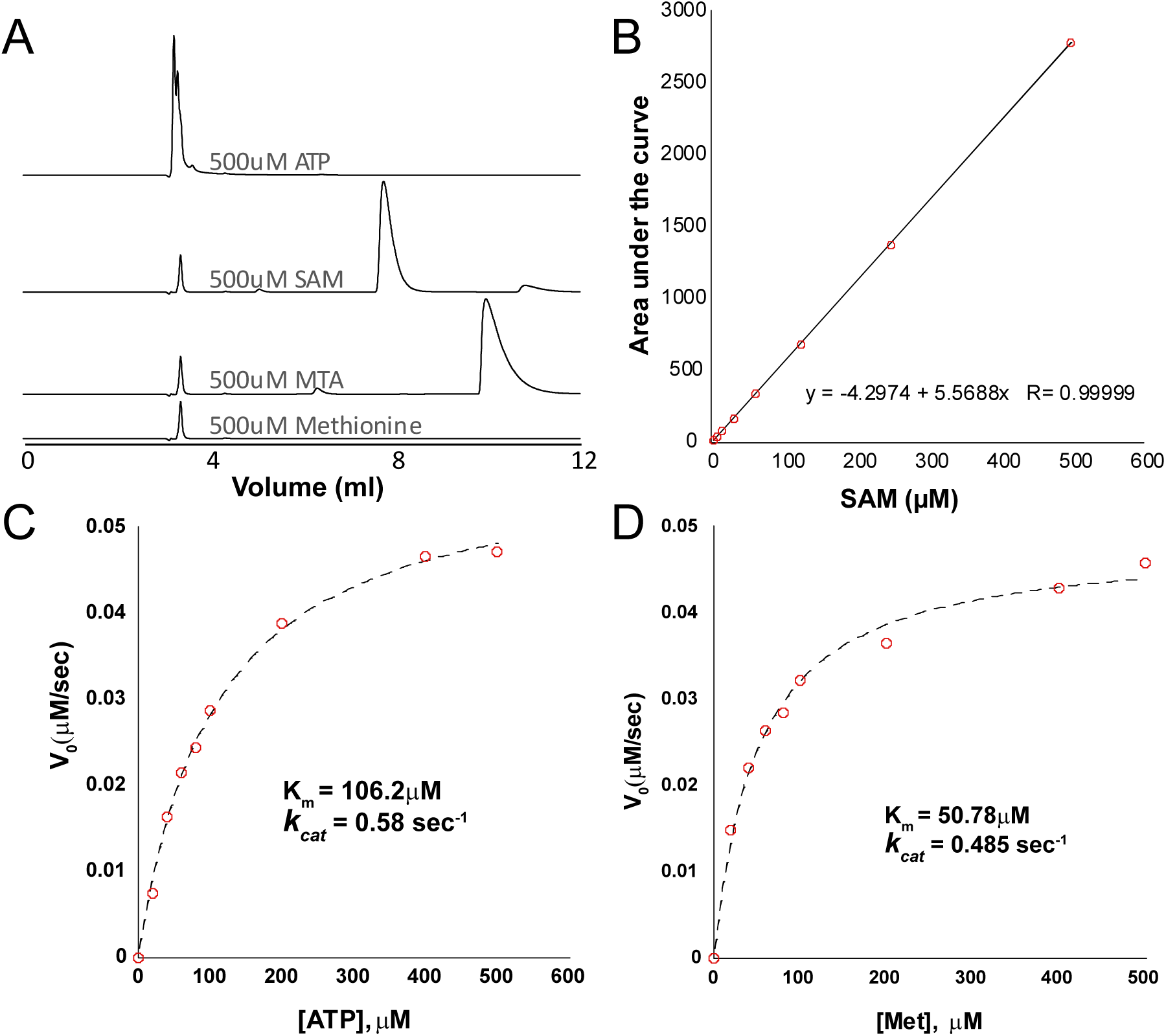
(A) ATP, SAM, methionine, and 5’-methylthioadenosine (MTA) standards separated on strong cation exchange column in HPLC instrument (see **STAR Methods**). (B) The integrated area of SAM traces produced by HPLC separation strongly correlates with the concentration of the injected SAM standards. (C) Michaelis-Menten fit of uMAT activity at a saturated amount of methionine and 0 to 500 μM ATP. (D) Michaelis-Menten fit of uMAT activity at a saturated amount of ATP and 0 to 500 μM methionine (see **STAR Methods**).

**Figure S5. Related to Figure 3.**
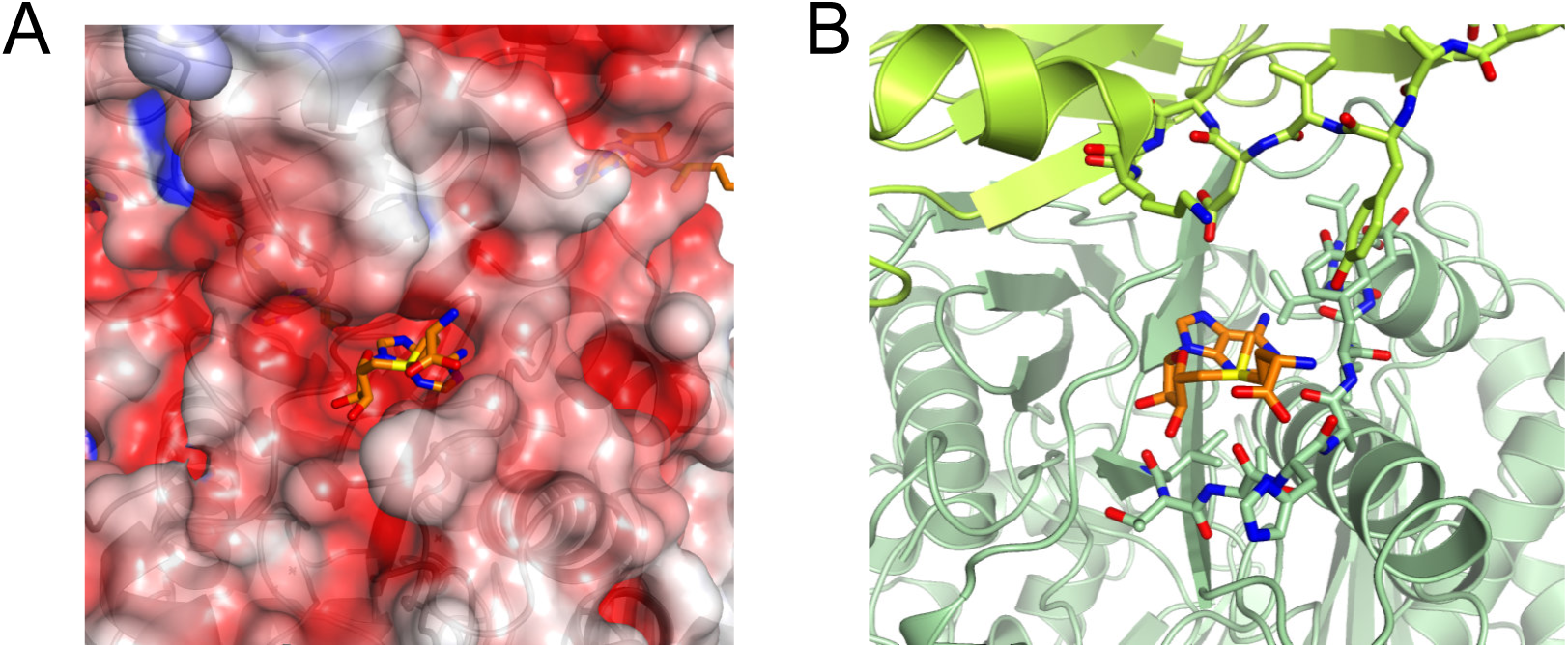
(A) Electrostatic surface representation reveals a negatively charged inter-dimeric pocket of uMAT (PDB ID: 6RK7) that accommodates a SAM molecule. (B) SAM molecule from the inter-dimeric interface of uMAT (PDB ID: 6RK7) is placed in the inter-dimeric interface of uMAT structure solved in the absence of ligands (PDB ID: 6RJS). Note that no steric hindrance exists that could have prevented SAM binding to the interface prior to the structural realignment of the dimers.

**Figure S6. Related to Figure 6.**
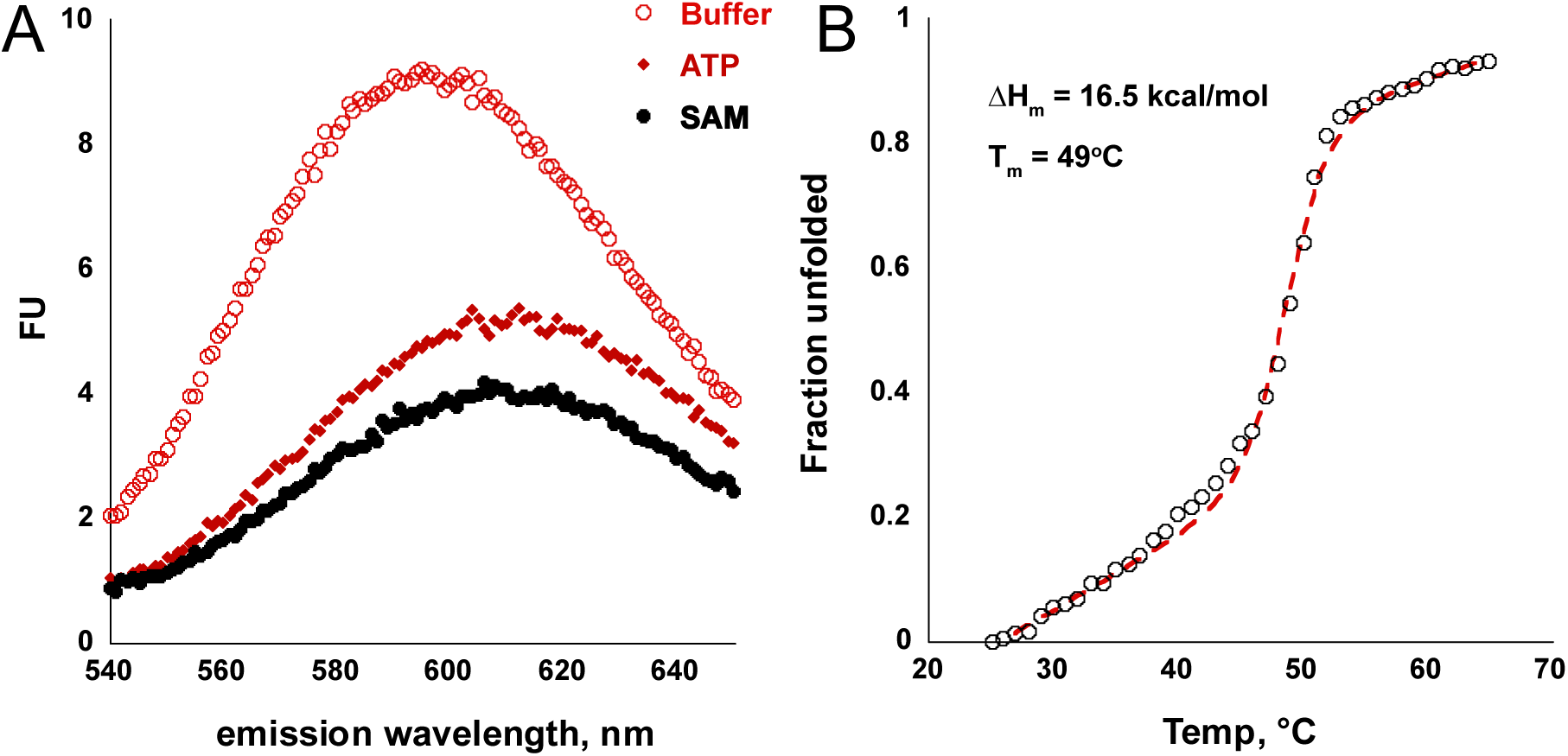
(A) uMAT SYPRO-Orange fluorescence signal in the absence (red open circles), and presence of ATP (red diamonds), and SAM (closed black circles). (B) Thermal unfolding of uMAT monitored by tryptophan fluorescence and fitted to two-state model. See **STAR Methods** for details.

**Figure S7. Related to Figure 7.**
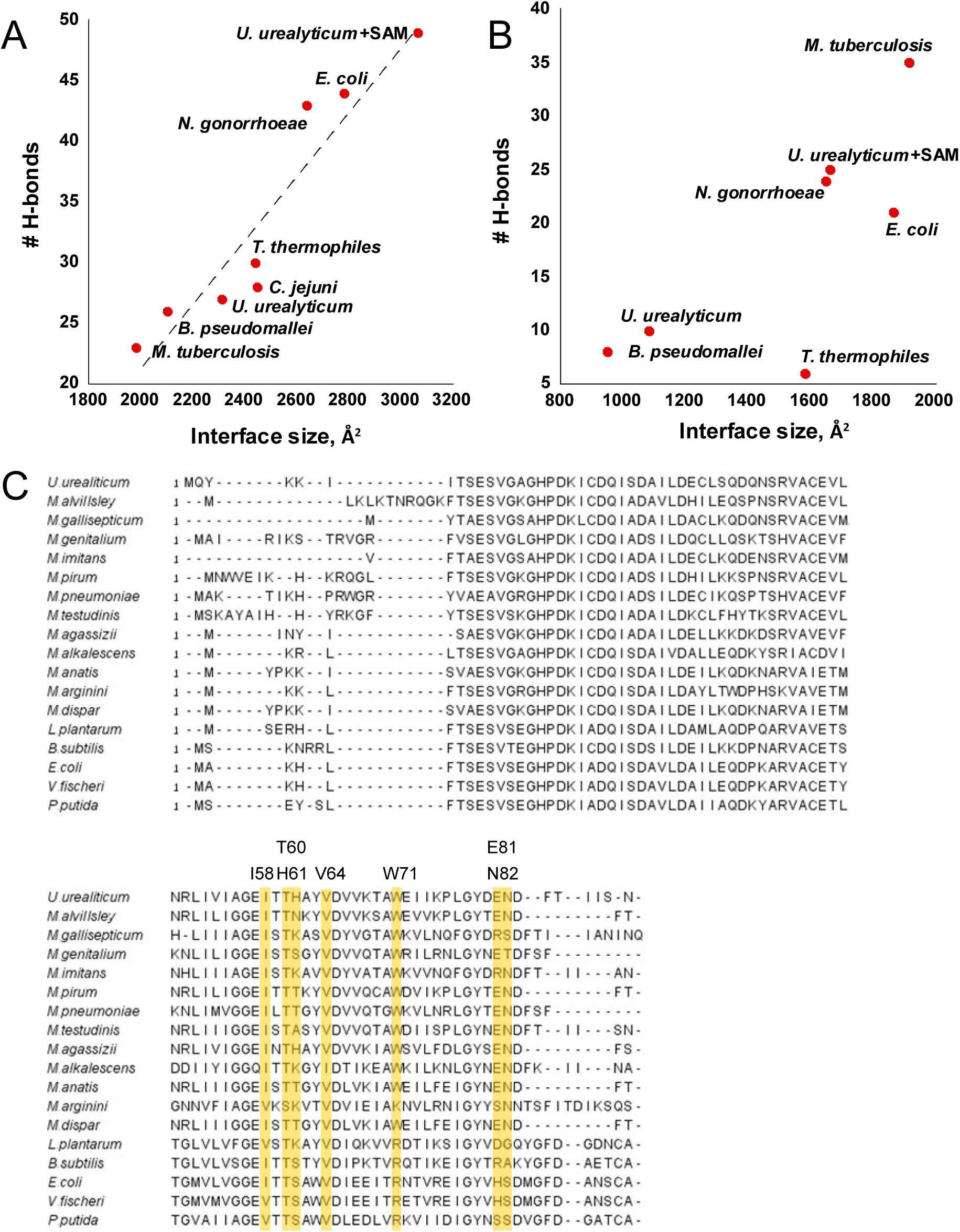
The correlation between the size of the interfaces and number of H-bonds in the large dimeric (A) and small inter-dimeric (B) interfaces, as detected by PISA (see Table 1, Table S2, and **STAR Methods**). (C) Multiple sequence alignment of bacterial MATs. Residues in the inter-dimeric interface forming direct interactions with SAM are highlighted in yellow and numbered according to uMAT sequence.

**Table S1.**
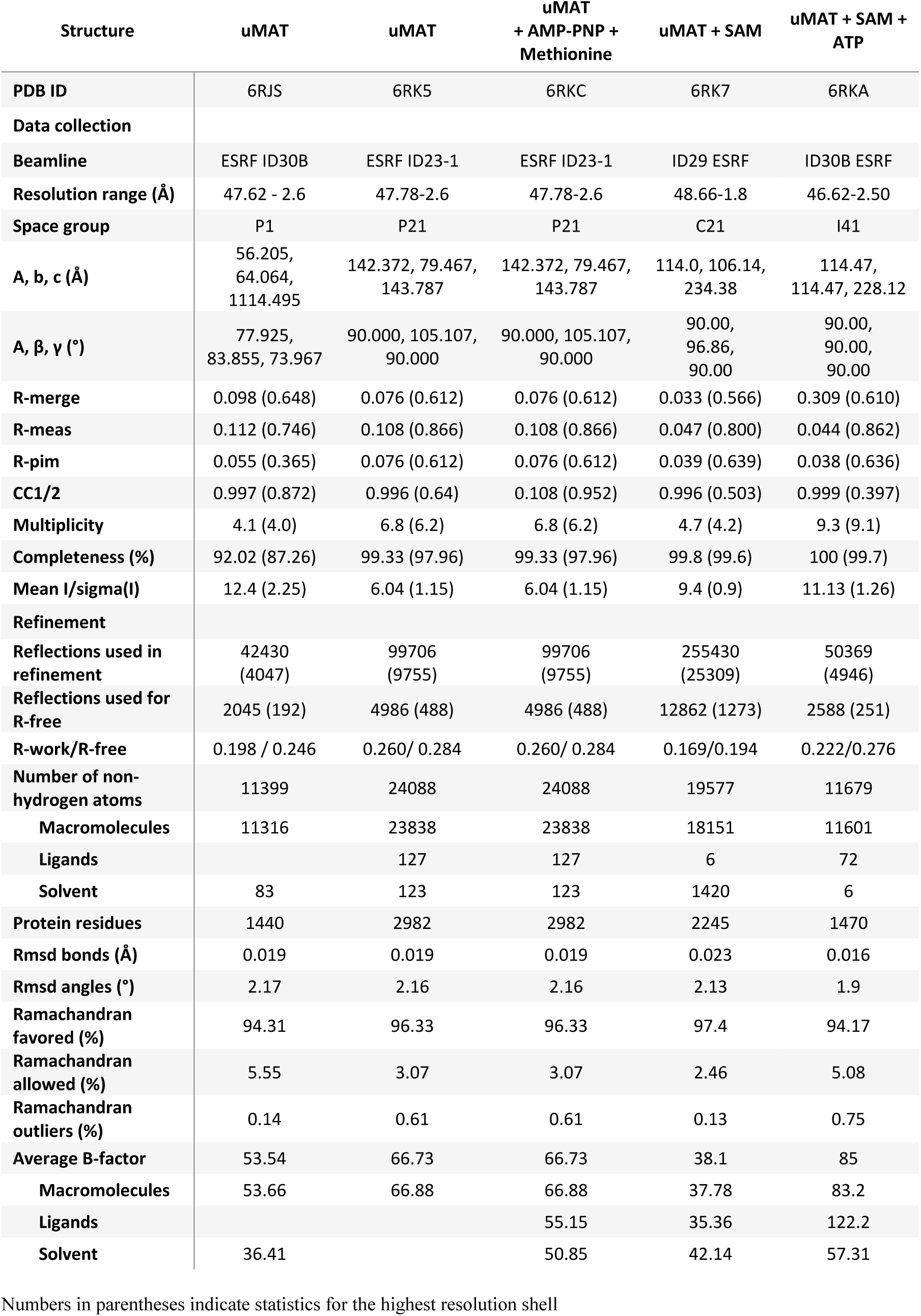
Crystallographic data collection and refinement statistics. Related to Figure 1 and Figure S1.

**Table S2.**
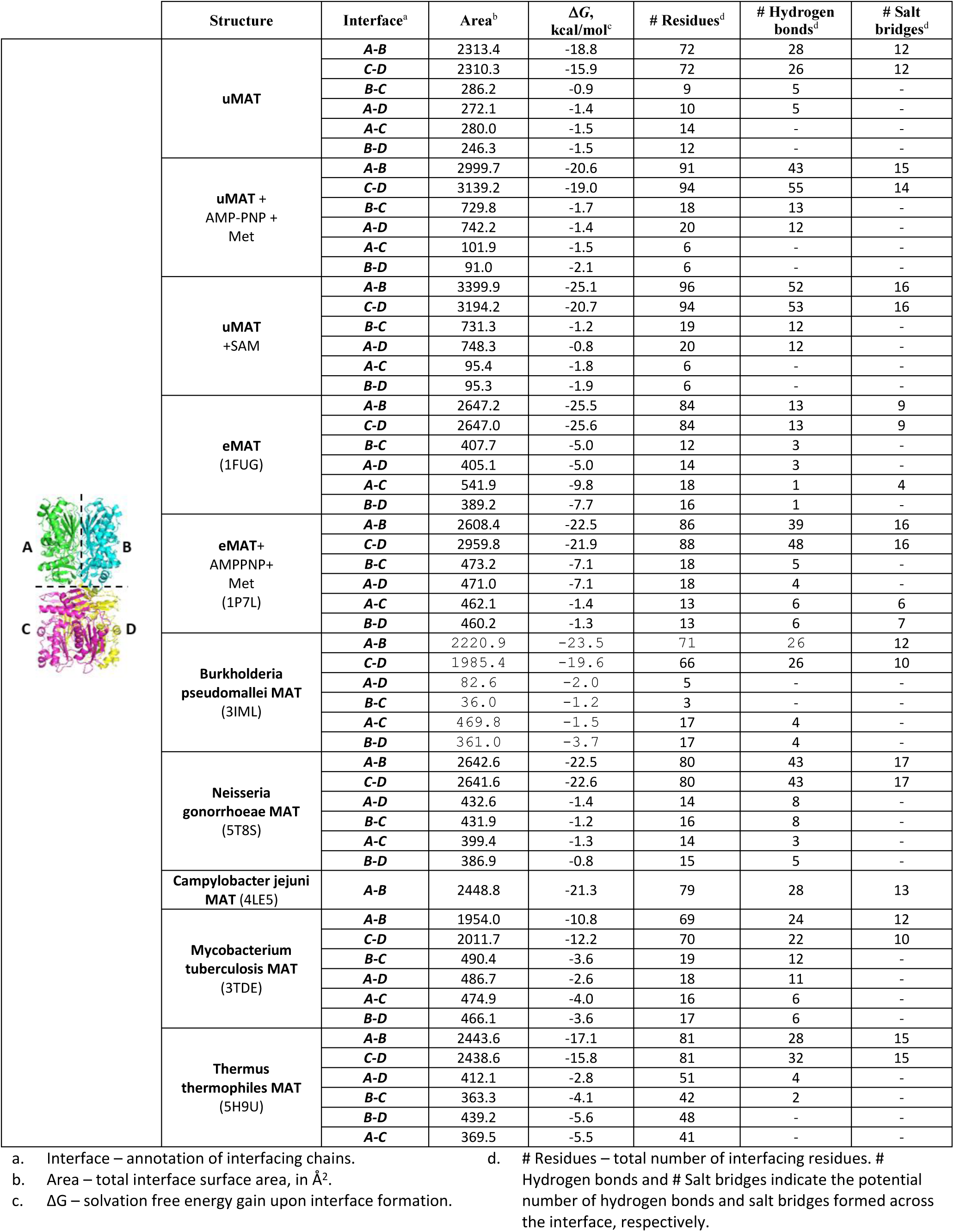
Extended interface size and composition analysis of orthologous MATs. Related to Table 1.

| | Structure | Interface<sup>a</sup> | Area<sup>b</sup> | $\Delta G$,<br>kcal/mol<sup>c</sup> | # Residues<sup>d</sup> | # Hydrogen<br>bonds<sup>d</sup> | # Salt<br>bridges<sup>d</sup> |
| --- | --- | --- | --- | --- | --- | --- | --- |
|  | uMAT | A-B | 2313.4 | -18.8 | 72 | 28 | 12 |
|  |  | C-D | 2310.3 | -15.9 | 72 | 26 | 12 |
|  |  | B-C | 286.2 | -0.9 | 9 | 5 | - |
|  |  | A-D | 272.1 | -1.4 | 10 | 5 | - |
|  |  | A-C | 280.0 | -1.5 | 14 | - | - |
|  |  | B-D | 246.3 | -1.5 | 12 | - | - |
|  | uMAT +<br>AMP-PNP +<br>Met | A-B | 2999.7 | -20.6 | 91 | 43 | 15 |
|  |  | C-D | 3139.2 | -19.0 | 94 | 55 | 14 |
|  |  | B-C | 729.8 | -1.7 | 18 | 13 | - |
|  |  | A-D | 742.2 | -1.4 | 20 | 12 | - |
|  |  | A-C | 101.9 | -1.5 | 6 | - | - |
|  |  | B-D | 91.0 | -2.1 | 6 | - | - |
|  | uMAT<br>+SAM | A-B | 3399.9 | -25.1 | 96 | 52 | 16 |
|  |  | C-D | 3194.2 | -20.7 | 94 | 53 | 16 |
|  |  | B-C | 731.3 | -1.2 | 19 | 12 | - |
|  |  | A-D | 748.3 | -0.8 | 20 | 12 | - |
|  |  | A-C | 95.4 | -1.8 | 6 | - | - |
|  |  | B-D | 95.3 | -1.9 | 6 | - | - |
|  | eMAT<br>(1FUG) | A-B | 2647.2 | -25.5 | 84 | 13 | 9 |
|  |  | C-D | 2647.0 | -25.6 | 84 | 13 | 9 |
|  |  | B-C | 407.7 | -5.0 | 12 | 3 | - |
|  |  | A-D | 405.1 | -5.0 | 14 | 3 | - |
|  |  | A-C | 541.9 | -9.8 | 18 | 1 | 4 |
|  |  | B-D | 389.2 | -7.7 | 16 | 1 | - |
|  | eMAT+<br>AMPPNP+<br>Met<br>(1P7L) | A-B | 2608.4 | -22.5 | 86 | 39 | 16 |
|  |  | C-D | 2959.8 | -21.9 | 88 | 48 | 16 |
|  |  | B-C | 473.2 | -7.1 | 18 | 5 | - |
|  |  | A-D | 471.0 | -7.1 | 18 | 4 | - |
|  |  | A-C | 462.1 | -1.4 | 13 | 6 | 6 |
|  |  | B-D | 460.2 | -1.3 | 13 | 6 | 7 |
|  | Burkholderia<br>pseudomallei MAT<br>(3IML) | A-B | 2220.9 | -23.5 | 71 | 26 | 12 |
|  |  | C-D | 1985.4 | -19.6 | 66 | 26 | 10 |
|  |  | A-D | 82.6 | -2.0 | 5 | - | - |
|  |  | B-C | 36.0 | -1.2 | 3 | - | - |
|  |  | A-C | 469.8 | -1.5 | 17 | 4 | - |
|  |  | B-D | 361.0 | -3.7 | 17 | 4 | - |
|  | Neisseria<br>gonorrhoeae MAT<br>(5T8S) | A-B | 2642.6 | -22.5 | 80 | 43 | 17 |
|  |  | C-D | 2641.6 | -22.6 | 80 | 43 | 17 |
|  |  | A-D | 432.6 | -1.4 | 14 | 8 | - |
|  |  | B-C | 431.9 | -1.2 | 16 | 8 | - |
|  |  | A-C | 399.4 | -1.3 | 14 | 3 | - |
|  |  | B-D | 386.9 | -0.8 | 15 | 5 | - |
|  | Campylobacter jejuni<br>MAT (4LE5) | A-B | 2448.8 | -21.3 | 79 | 28 | 13 |
|  | Mycobacterium<br>tuberculosis MAT<br>(3TDE) | A-B | 1954.0 | -10.8 | 69 | 24 | 12 |
|  |  | C-D | 2011.7 | -12.2 | 70 | 22 | 10 |
|  |  | B-C | 490.4 | -3.6 | 19 | 12 | - |
|  |  | A-D | 486.7 | -2.6 | 18 | 11 | - |
|  |  | A-C | 474.9 | -4.0 | 16 | 6 | - |
|  |  | B-D | 466.1 | -3.6 | 17 | 6 | - |
|  | Thermus<br>thermophilus MAT<br>(5H9U) | A-B | 2443.6 | -17.1 | 81 | 28 | 15 |
|  |  | C-D | 2438.6 | -15.8 | 81 | 32 | 15 |
|  |  | A-D | 412.1 | -2.8 | 51 | 4 | - |
|  |  | B-C | 363.3 | -4.1 | 42 | 2 | - |
|  |  | B-D | 439.2 | -5.6 | 48 | - | - |
|  |  | A-C | 369.5 | -5.5 | 41 | - | - |

- a. Interface – annotation of interfacing chains. b. Area – total interface surface area, in Å<sup>2</sup>. c. $\Delta G$ – solvation free energy gain upon interface formation.
- d. # Residues – total number of interfacing residues. # Hydrogen bonds and # Salt bridges indicate the potential number of hydrogen bonds and salt bridges formed across the interface, respectively.

